# Compartmentalized lymph node drainage dictates intestinal adaptive immune responses

**DOI:** 10.1101/299628

**Authors:** Daria Esterházy, Maria CC Canesso, Paul A Muller, Ainsley Lockhart, Luka Mesin, Ana MC Faria, Daniel Mucida

## Abstract

The intestinal immune system has the challenging task of tolerating foreign nutrients and the commensal microbiome, while excluding or eliminating ingested pathogens. Failure in such balance leads to a range of severe intestinal and systemic diseases such as inflammatory bowel diseases, food allergies or invasive gastrointestinal infections^1,2^. Multiple innate and adaptive immune mechanisms are therefore in place to maintain tissue integrity, including efficient peripheral generation of effector T (T_H_) cells and FOXP3^+^ regulatory T (pTreg) cells, which mediate resistance to pathogens and regulate excessive immune activation, respectively^2–5^. The gut-draining mesenteric lymph nodes (mLNs) are critical sites for orchestrating adaptive immunity to luminal perturbations^6–8^. However, how they manage to simultaneously support tolerogenic and inflammatory reactions is incompletely understood. Here we report that individual mLNs are anatomically and immunologically distinct according to the functional gut segment they drain. Dendritic cell gene signatures and adaptive T cell polarization against the same luminal antigen differed between mLNs along the intestine, the proximal small intestine–draining mLNs preferentially giving rise to tolerogenic and the distal mLNs to pro-inflammatory T cell responses. This compartmentalized dichotomy could be perturbed by duodenal infection, surgical removal of select distal mLNs, dysbiosis, or ectopic antigen delivery, impacting both lymphoid organ and tissue immune responses. Our findings reveal that the conflict between tolerogenic and inflammatory adaptive responses is in part resolved by discrete mLN drainage, and encourage gut segment-specific antigen targeting for therapeutic immune modulation.

---

The intestine’s main function is to digest and absorb dietary nutrients, with help from the absorptive epithelium and underlying vasculature and lymphatic system, as well as the microbiome^6^. Nutrient selection and absorptive capacity change along the intestine; most dietary nutrients are absorbed in the villi of the upper small intestine (duodenum (D) and jejunum (J)) and to a lesser extent by the lower small intestine (ileum, I), while the dense microbiota in the I and colon (C) mediates further nutrient release. The intestine also houses the largest immune compartment in the body, tasked with providing resistance to toxins and invading pathogens while maintaining tolerance to dietary and microbiota antigens, either by local action or lymphatic trafficking to the gut-draining mLNs to mount adaptive responses^1,7,9^. We sought to understand how compartmentalized lymphatic drainage of the intestine contributes to an organized immune response in this complex environment.

To address the role of tissue drainage for immune responses triggered along the intestine, we first imaged the gut lymphatic system including the intestine, mesentery and mLNs using 3D imaging of solvent-cleared tissue stained with an antibody against the lymphatic endothelial cell (LEC) surface marker LYVE-1. In the gut its structure followed the proximal to distal intestinal gradient of absorptive capacity^10^, whereby lacteal length decreased along the small intestine (Fig. 1 a-c). In germ-free (GF) mice, duodenal absorption may compensate for lack of microbiome-mediated distal nutrient release^11^, and this was reflected in the length of lymphatic lacteals (Fig. 1c). In contrast, the lymphatics of the colon are organized in two distinct layers, lack lacteals and did not appear to differ in GF mice, suggesting that its structural organization is independent of the microbiome (Fig. 1c). The cleared-tissue visualization also revealed a seemingly continuous submucosal lymphatic network along the intestine (Fig. 1a-c, Extended Data Fig. 1a, Supplementary Video 1-3). The local gut lymphatics are linked via afferent lymphatics in the mesentery to the mLNs, which thus provide a physical route for immune cell trafficking including CD11c^+^ antigen-presenting dendritic cells (DCs) and may maintain a region-specific compartmentalization (Fig. 1d-h). Our high-resolution analysis confirmed that in both specific pathogen-free (SPF) and GF mice, collecting vessels from different gut segments connected to distinct mLNs and were densely packed around the nodes suggesting high efflux of lymph (Fig. 1g, h, Extended Data Fig. 1b-e, Supplementary videos 4-7). To substantiate that gut segment-specific lymphatic drainage leads to compartmentalized mLN uptake, we took advantage of the fact that dietary lipids are packaged into chylomicrons in duodenal epithelium and are preferentially absorbed via the lacteals of the upper small intestine. We tracked radiolabelled retinol, as a lipid soluble proxy and an immuno-modulatory nutrient^12^, following intra-gastric delivery. Indeed, most retinol was absorbed in the duodenum, and this was mirrored in the D-mLNs. This was dependent on the lymphatic route, as treatment with a chylomicron formation inhibitor prevented duodenal retinol efflux to its draining mLN (Fig. 1i-k, Extended Data Fig. 1f, g). Similarly, the increased duodenal but decreased ileal absorptive capacity in GF mice was displayed by corresponding retinol retention in gut tissue and mLNs (Extended Data Fig. 1h-j). These data suggest that the compartmentalized lymphatic route transmits distinct niches along the intestine to discrete secondary lymphoid organs, potentially contributing to a regionally-organized immune response.

**Figure 1.**
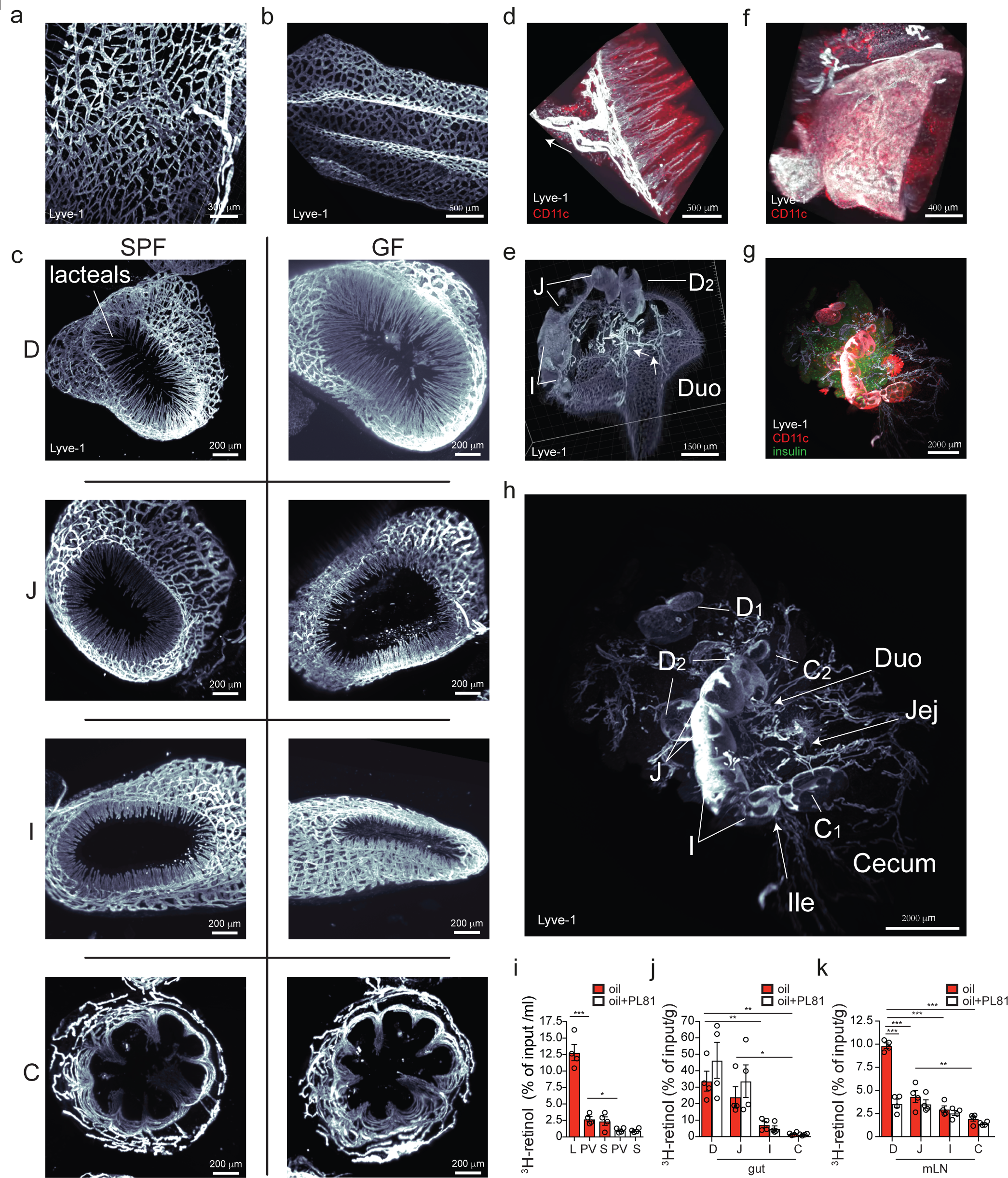
The intestinal lymphatics provide a route for shaping distinct mLN environments. **a-h**, 3D reconstruction of mouse lymphatics (α-LYVE-1) after solvent clearing (iDISCO+) and light-sheet microscopy of the duodenal (**a**) and colonic (**b**) submucosa, villi and submucosa along the intestine (**c**), duodenal mesenteric collecting vessels (**d**), the central chain of mLNs (**e**), the colonic mLN (**f**), all mLNs and afferent lymphatics coming from the gut (**g**, **h**). The following specifics apply: **c**, 9 weeks old GF and SPF mice, lymphatics protruding into the villi (lacteals) are pointed out; **d**, **e**, arrows denote direction of lymph flow from intestine to mLNs; **d**, **f**, **g**, CD11c was revealed by using Itgax-Venus mice and staining against GFP; **g**, α-insulin was used to indicate the mLNs embedded in the pancreas. D/Duo=duodenum, whereby D1 portal LNs and D2 distal duodenum mLNs, J/Jej=jejunum, I/Ile=ileum, C1=cecal-colonic mLN, C2=ascending colonic mLN. **i-k**, Percentage of ^3^H-retinol absorption into lymph (L), portal vein serum (PV) or systemic serum (S) (**i**), indicated gut segment (**j**) or mLN (**k**) of mice 3 h after gavage with 1 µCi ^3^H-retinol in 100 µl olive oil with or without pre-treatment with 5 µl chylomicron formation inhibitor Pluronic L-81 3h prior to gavage (*n*= 4 per group). \**P* < 0.05, \*\**P* < 0.01 and \*\*\**P* < 0.005 (one-tailed *t*-test or ANOVA). Data representative of two independent experiments.

**Extended Figure 1.**
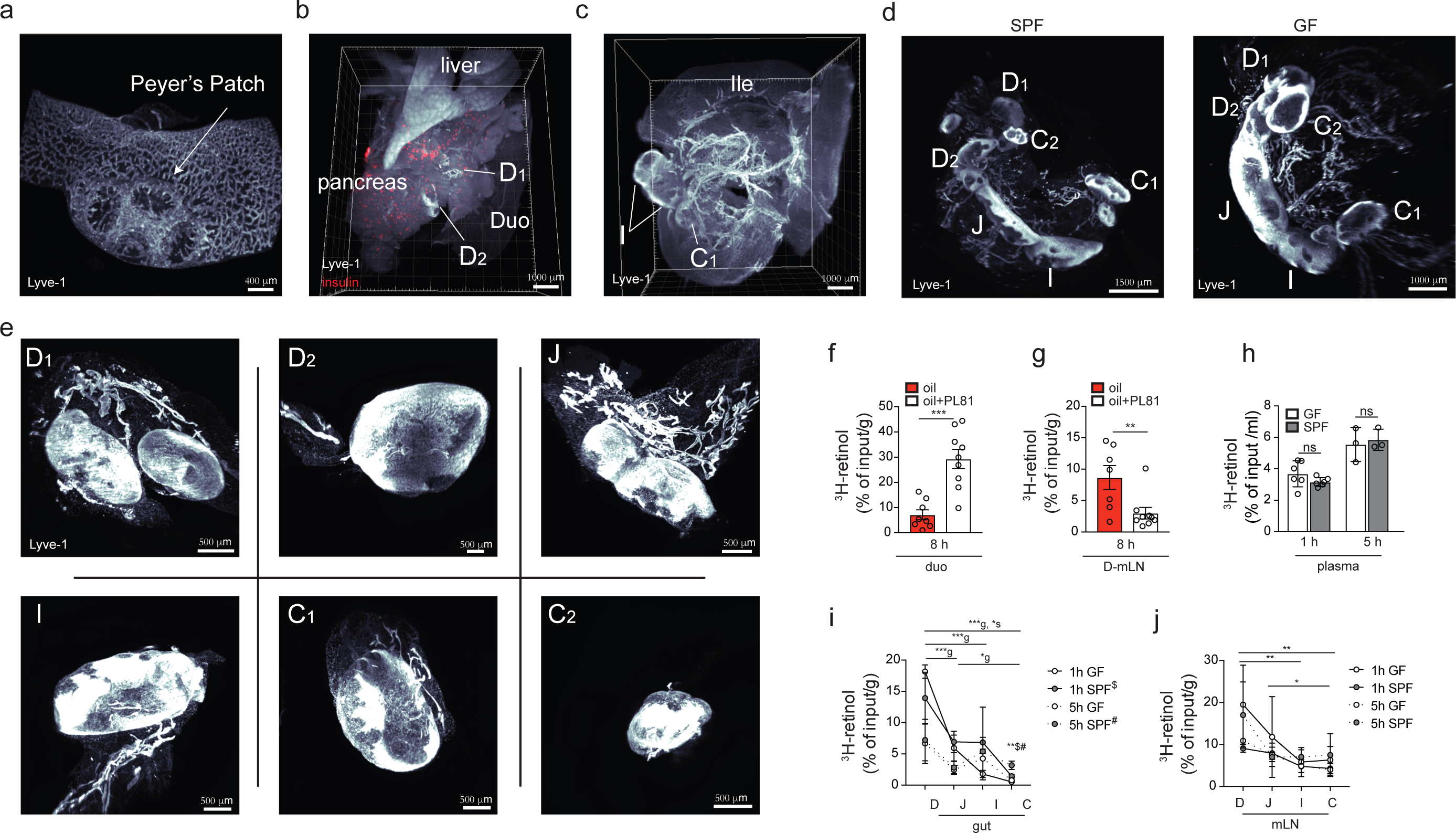
Extended characterization of the organization of intestinal immune system lymphatics and dietary retinol absorption. **a-e**, 3D reconstruction of mouse lymphatics (α-LYVE-1) after solvent clearing (iDISCO+) and light sheet microscopy of an ileal Peyer’s Patch (**a**), the duodenal (**b**) and distal intestine (**c**) mLNs, the entire mLN chain of SPF and GF mice (**d**), and the individually dissected mLNs (**e**). The following specifics apply: **b**, α-insulin was used to indicate the duodenal mLNs embedded in the pancreas, and liver and pancreas are indicated**; d**, 9 weeks old GF and SPF mice. D/Duo=duodenum, whereby D1 portal LNs and D2 distal duodenum mLNs, J/Jej=jejunum, I/Ile=ileum, C1=cecal-colonic mLN, C2=ascending colonic mLN. **f**, **g**, Percentage of ^3^H-retinol absorption into duodenum (**f**), or duodenal mLN (**g**) of mice 8 h after gavage with 1 µCi ^3^H-retinol in 100 µl olive oil with or without pre-treatment with 5 µl chylomicron formation inhibitor Pluronic L-81 3h prior to gavage (*n*= 8-9 per group). Data pooled from two independent experiments with 4-5 animals per group each. **h**-**j**, Percentage of ^3^H-retinol absorption into systemic plasma (**h**), indicated intestinal tissue (**i**) or mLN (**j**) of GF or SPF mice 1 or 5 h after gavage with 1 µCi ^3^H-retinol in 100 µl olive oil (*n*= 3 per group). Data representative of two independent experiments. \**P* < 0.05, \*\**P* < 0.01 and \*\*\**P* < 0.005 (one-tailed *t*-test or ANOVA). g refers to GF ANOVA, s to SPF ANOVA, $ to 1 h time point, # to 5 h time point.

We asked if gut segment-specific lymphatic drainage influences immune cells in the mLNs, particularly migratory DCs, which are known to initiate tissue-specific immune responses^13^. The two MHCII^hi^ migratory DC populations, CD103^+^CD11b^−^ and CD103^+^CD11b^+^ DCs, are highly represented in mLNs and have been shown to support tolerogenic and pro-inflammatory responses, respectively^13, 14^. We sorted these two populations from D, I and C-draining mLNs and performed RNA-seq. Principal component analysis revealed that DCs segregated strongly between small versus large intestine and less between D and I; these effects were more pronounced amongst the CD103^+^CD11b^+^ than CD103^+^CD11b^−^ DCs (Fig. 2a-c, Extended Data Fig. 2a-d), possibly reflective of the fact that the former are the more abundant population in the intestinal lamina propria^14^ (Extended Data Fig. 2e-i). The stronger impact of the environment on CD103^+^CD11b^+^ DCs was also evident in the differential gene expression and gene set enrichment analysis of cellular pathways, though both migratory DC subsets exhibited immunological and metabolic differences by mLN location (Fig. 2b, c, Extended Data Fig. 2a-d). Most notably, both D-mLN populations displayed a less pro-inflammatory signature than their colonic counterparts: D-mLN CD103^+^CD11b^+^ DCs expressed lower levels of inflammatory cytokine receptors and pathways such as interferon or IL-1β (Fig. 2b); D-mLN CD103^+^CD11b^−^ DCs were specifically enriched for *Ccl22*, encoding a chemokine associated with migration of Tregs^15^, and Treg-promoting factor *Aldh1a2*, a rate-limiting enzyme for retinoic acid (RA)^12,16^ production from dietary retinol (Fig. 2c, d). An enzymatic activity assay confirmed that RA production capacity of DCs was overall higher in the small than in the large intestine-draining mLN DCs, and preserved in CD103^+^CD11b^−^ DCs from GF mice (Fig. 2d, Extended Data Fig. 2e), albeit the ratio of tolerogenic CD103^+^CD11b^−^ DCs to pro-inflammatory CD103^+^CD11b^+^DCs^7,14^ was inverted in all mLNs in the absence of a microbiota (Fig. 2e). These findings suggest the D-mLNs as tolerogenic environments due to favourable DC signatures and nutrient drainage, a phenomenon boosted by the microbiota.

**Figure 2.**
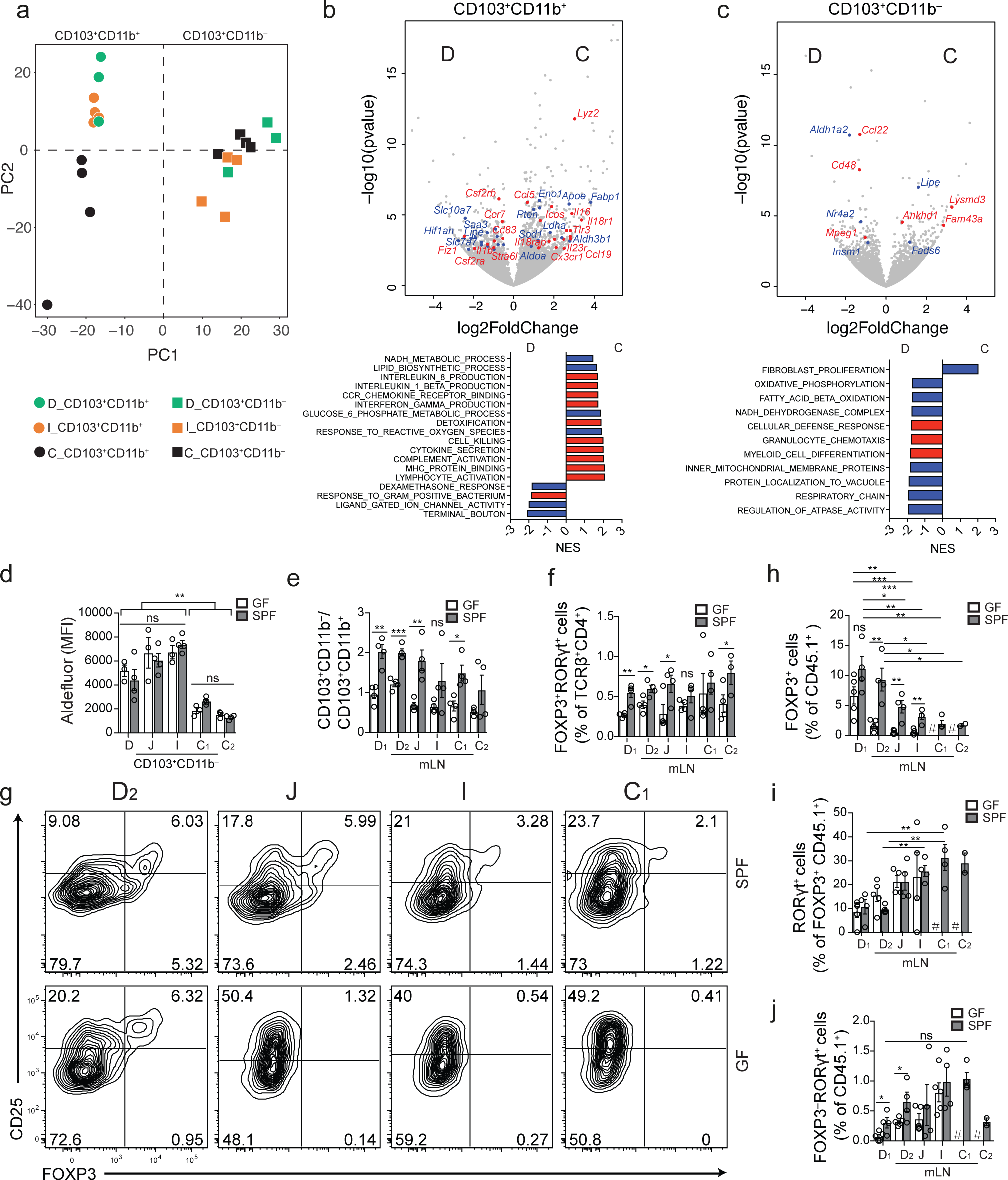
Dendritic cell properties and CD4^+^ T cell polarization by dietary antigen differ along the mLN chain. **a**, Principle component analysis of RNA-seq of CD103^+^CD11b^+^ and CD103^+^CD11b^−^ DCs sorted from the duodenal (D), ileal (I) or cecal-colonic (C) mLN of 15 weeks old B6 mice reared under SPF conditions (*n*=3-4, pooled from 2 mice each). PC1 accounts for 23%, PC2 for 10% of the variance. **b**, **c**, Volcano plot depicting differentially expressed genes (top) and bar graph showing some differentially regulated pathways (bottom) among CD103^+^CD11b^+^ (**b**) and CD103^+^CD11b^−^ (**c**) DCs from the duodenal (D) or cecal-colonic (C) mLNs identified by RNA-seq. Blue dots and bars indicate metabolic genes and pathways, red dots and bars immunity-related genes and pathways. **d**, Mean fluorescence intensity (MFI) of fluorescein isothiocyanate-positive boron-dipyrromethene-tagged aminoacetate, the ALDH product of boron-dipyrromethene-tagged aminoacetaldehyde (non-toxic substrate for ALDH) (Aldefluor) in CD103^+^CD11b^−^ DCs isolated from indicated mLNs of GF and SPF mice (*n*=3), assessed by flow cytometry 30 min after the addition of substrate. **e**, Ratio of CD103^+^CD11b^+^ and CD103^+^CD11b^−^ DCs in indicated mLNs of 7 weeks old SPF and GF mice (*n*=4) as determined by flow cytometry. Data representative of two independent experiments. **f**, Frequency of FOXP3^+^RORγt^+^ among TCRβ^+^CD4^+^ cells in indicated mLN of GF or SPF mice (*n*=3). Data representative of two independent experiments. **g**, Representative flow cytometry plots of Treg (FOXP3^+^) and activated cell (CD25^+^) frequency in indicated mLNs of GF and SPF mice. **h**-**j**, Frequency of total FOXP3^+^ among CD45.1^+^ (**h**), RORγt^+^ among FOXP3^+^ CD45.1^+^ (**i**), and RORγt^+^ among CD45.1^+^ (**j**) cells in indicated mLNs 60 h post adoptive transfer of 1 x 10^6^ naïve CD45.1^+^ OT-II cells into GF and SPF CD45.2 hosts (*n*=4) and after gavage of OVA 48 h and 24 h prior to analysis. Data representative of two independent experiments. # indicates fewer than 200 cells were recovered. D=duodenum, whereby D1 portal LNs and D2 distal duodenum mLNs, J=jejunum, I=ileum, C1=cecal-colonic mLN, C2=ascending colonic mLN, C=colon. \**P* < 0.05, \*\**P* < 0.01 and \*\*\**P* < 0.005 (one-tailed *t*-test or ANOVA).

**Extended Figure 2.**
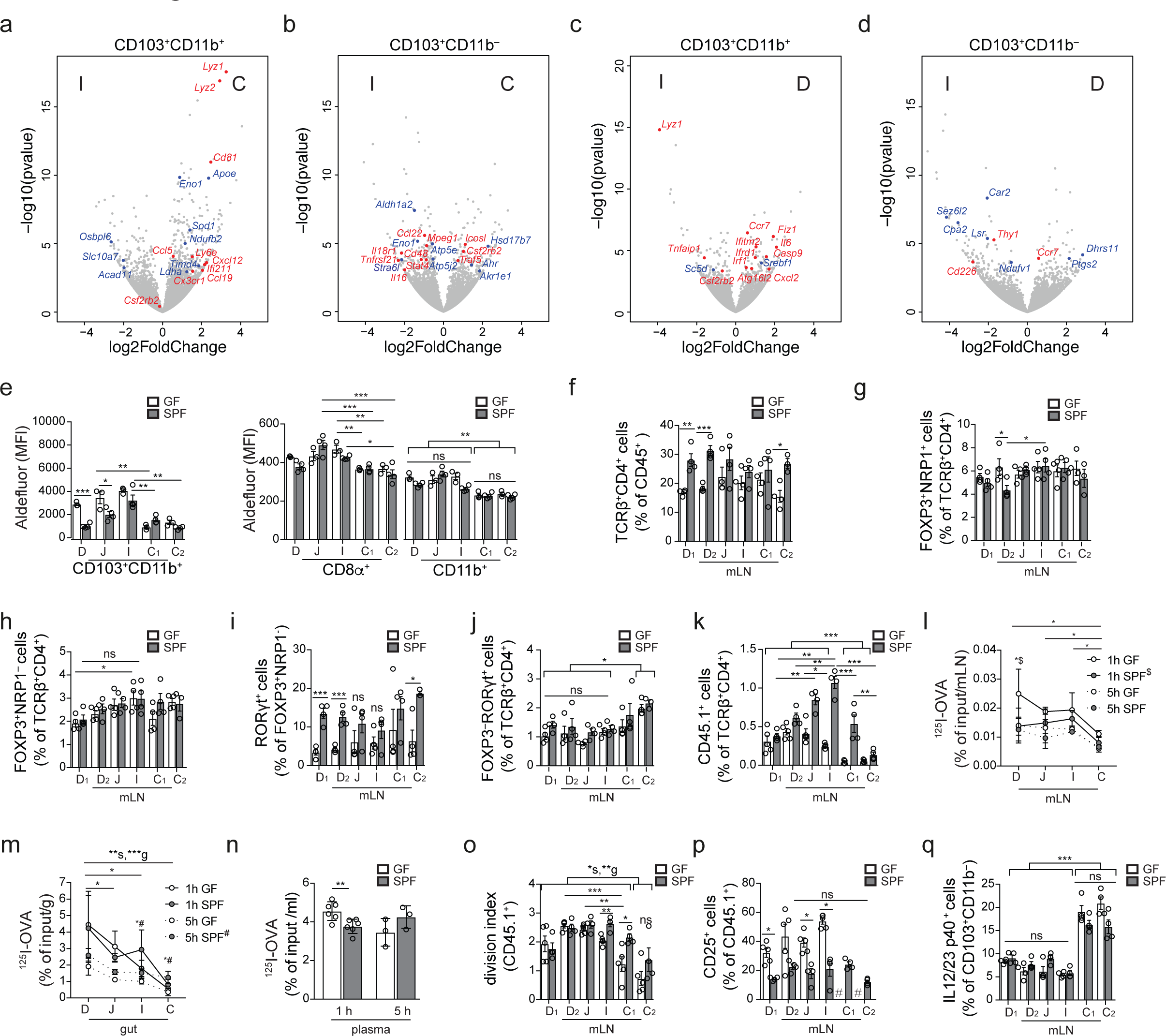
Extended analysis of immune cell populations and dietary antigen absorption in different mLNs. **a**-**d**, Volcano plot depicting differentially expressed genes among CD103^+^CD11b^+^ and CD103^+^CD11b^−^ DCs sorted from ileal versus colonic mLNs (**a**, **b**) or ileal versus duodenal mLNs (**c**, **d**) identified by RNA-seq. Blue dots indicate metabolic, red dots immunity-related genes. **e**, Mean fluorescence intensity (MFI) of fluorescein isothiocyanate-positive boron-dipyrromethene-tagged aminoacetate, the ALDH product of boron-dipyrromethene-tagged aminoacetaldehyde (non-toxic substrate for ALDH) (Aldefluor) in indicated DC populations and mLNs from GF and SPF mice (*n*=3) assessed by flow cytometry 30 min after the addition of substrate. **f**-**j**, Frequency of TCRβ^+^CD4^+^ among CD45^+^ (**f**), FOXP3^+^NRP1^+^ (**g**) and FOXP3^+^NRP1^−^ (**h**) among TCRβ^+^CD4^+^, RORγt^+^ among FOXP3^+^NRP1^−^ (**i**) and RORγt^+^ among TCRβ^+^CD4^+^ (**j**) cells in indicated mLN of GF or SPF mice (*n*=4). Data representative of two independent experiments. **k**, Frequency of CD45.1^+^ among TCRβ^+^CD4^+^ cells in indicated mLNs 60 h post adoptive transfer of 1 x 10^6^ naïve CD45.1^+^ OT-II cells into GF and SPF CD45.2 hosts (*n*=4) and after gavage of OVA 48 h and 24 h prior to analysis. **l-n**, ^125^I recovery in indicated mLNs (**l**), intestinal tissue (**m**) and plasma (**n**) of GF and SPF mice (*n*=3 per time point and group) 1h or 5 h after gavage with 4 x 10^6^ CPM ^125^I-OVA in 50 mg cold OVA. **o**, **p**, Division index of CD45.1^+^ (o) and frequency of CD25^+^ among CD45.1^+^ cells in indicated mLNs as in **k**. **q**, Frequency of IL-12/23 p40^+^ cells among CD103^+^CD11b^−^ DCs in indicated mLNs of GF and SPF mice (*n*=4 per group). # indicates fewer than 200 cells were recovered. D=duodenum, whereby D1 portal LNs and D2 distal duodenum mLNs, J=jejunum, I=ileum, C1=cecal-colonic mLN, C2=ascending colonic mLN, C=colon. \**P* < 0.05, \*\**P* < 0.01 and \*\*\**P* < 0.005 (one-tailed *t*-test or ANOVA).

Since migratory DCs play a pivotal role in instructing adaptive immune cells, we next investigated if CD4^+^ T cell fates also differed between mLNs and correlated with migratory-DC profiles or with the presence of the microbiota. Although GF mice harbour reduced Treg cells in the small and large intestines^17,18^, there was no difference in Treg cell frequency between mLNs of GF and SPF mice or along the mLN chain (Extended Data Fig. 2f-h). T_H_17 effector cells and their corresponding suppressive RORγt^+^ pTreg cells can be induced by the microbiota^2–4^, and indeed GF mice displayed a severe reduction in RORγt^+^ pTreg cells in all mLNs (Fig. 2f, Extended Data Fig. 2i), although we did not observe a significant decrease in RORγt^+^ (T_H_17) cells (Extended Data Fig. 2j). To directly address if initial CD4^+^ T cell polarization occurs in a compartmentalized manner in gut-draining LNs, we adoptively transferred naïve Ovalbumin (OVA)-specific CD4^+^ cells (CD45.1 OT-II cells) and analysed their fate 48 h after OVA exposure by gavage^14^. We observed that in GF mice only the D-mLNs retained OT-II cells equally to SPF mice, while OT-II frequencies were much lower in J and I and almost undetectable in C-draining mLNs (Extended Data Fig. 2k). This was paralleled by a decreased absorption of OVA into the distal gut and its draining mLNs in GF versus SPF mice (Extended Data Fig. 2l-n) and mirrored the altered lymphatic structure in GF mice (Fig. 1c), suggesting that a gradient of antigen availability dictated OT-II cell frequencies in mLNs. Amongst the retained OT-II cells, we observed a clear gradient of pTreg induction that declined in a proximal to distal manner (Fig. 2g, h). In GF mice, pTreg induction was decreased in all mLNs (Fig. 2g, h), but not due to lack of activation per se (Extended Data 2o, p), suggesting the less favourable DC composition in GF mLNs contributed to this effect. Limited antigen availability alone was also unlikely to explain the gradient of OT-II pTreg cells in SPF mice, as CD25^+^ frequencies did not differ between mLNs (Extended Data Fig. 2p), and rather suggested compartmentalized difference in lymphatic retinol absorption and DC expression profiles as underlying causes. In support of such a role of a local environment, OT-II RORγt^+^ T_H_ and RORγt^+^ pTreg cells displayed an ascending proximal to distal gradient that was independent of the microbiota (Fig. 2i, j), and correlated with an increase in IL12/23p40^+^ frequency amongst the tolerogenic DCs (Extended Data Fig. 2q). These data indicate that under homeostatic conditions the duodenal mLNs are the primary sites for FOXP3^+^ pTreg induction by dietary antigen, implying that this anatomical site may be instrumental in preventing food intolerance or allergy. In contrast, even in response to the same antigen, distal mLNs preferentially promote T_H_17 and RORγt^+^ pTreg differentiation, suggesting a cooperation of efficient effector and corresponding regulatory mechanisms.

The mLNs are crucial for both oral tolerance and pathogen-resistance mechanisms^8,19,20^. Viral and bacterial gastrointestinal infections were shown to perturb the generation of pTreg cells recognizing dietary antigen and subsequent oral tolerance^7,21^, but it remains unclear whether these effects are linked to the infected intestinal regions and the associated mLNs. To induce an immunological conflict in the proximal intestine-draining mLNs, we chose the helminth *Strongyloides venezuelensis* (S.v.) as a rodent model for human *Strongyloides* infection that displays distinct duodenal tropism. S.v. larvae infect via the skin, and intestinal adult worm load peaks at 6-8 days post infection (p.i.) and is cleared after 10-12 days^22^. We observed that upon S.v. infection only the D-mLNs were swollen and displayed increased immune cell counts compared to mLNs of non-infected (N.I.) mice (Fig. 3a, b, Extended Data Fig. 3a), in correlation with compartmentalized duodenal worm load (Extended Data Fig. 3b). We also observed a D-mLN selective influx of CD11b^+^ DCs, previously linked to the induction of Th2 responses^23^, at the expense of the more tolerogenic CD103^+^CD11b^−^ and CD8α^+^ DC subsets (Fig. 3 c-e and Extended Data Fig. 3c-e), as well as a type 2 immunity signature comprising eosinophils, GATA3^+^ T_H_2 cells and GATA3^+^ Treg cells (Fig. 3f-h, Extended Data Fig. 3f). Endogenous polyclonal GATA3^−^ Treg and CD25^+^ CD4^+^ T cell frequencies were unaffected (Extended Data Fig. 3g-i); however, transferred OT-II cells displayed a significant reduction in upregulation of FOXP3 and CD25 specifically in the D-mLNs upon OVA gavage compared to N.I. mice (Fig. 3i, j, Extended Data Fig. 3j), despite proportional mLN seeding (Extended Data Fig. 3k, l), suggesting that the tolerogenic environment had been perturbed enough to impact de novo Treg induction or adaptation against dietary antigen. Although chronic helminth infection is generally thought to suppress allergies^24,25^, we asked if transient infections such as with S.v. during the initial antigen exposure impacted tolerance to dietary antigens, a pTreg-dependent process^5,26^. We fed OVA at day 6 and 7 post-S.v. infection and subjected mice to the asthma model of oral tolerance post-pathogen clearance (Fig. 3k). Oral tolerance is a very robust mechanism^14^, but nonetheless we observed a partial impairment of suppression of eosinophilia and DC infiltration into the bronchial-alveolar fluid (BALF) and lungs (Fig. 3 l, m, Extended Data Fig. 3m, n), and increased OVA-specific IgG1 levels in OVA-fed S.v. group compared to N.I. mice (Fig. 3n). Taken together these data show that duodenal infection can perturb local mLN responses as well as impact systemic tolerance to gut antigens.

**Figure 3.**
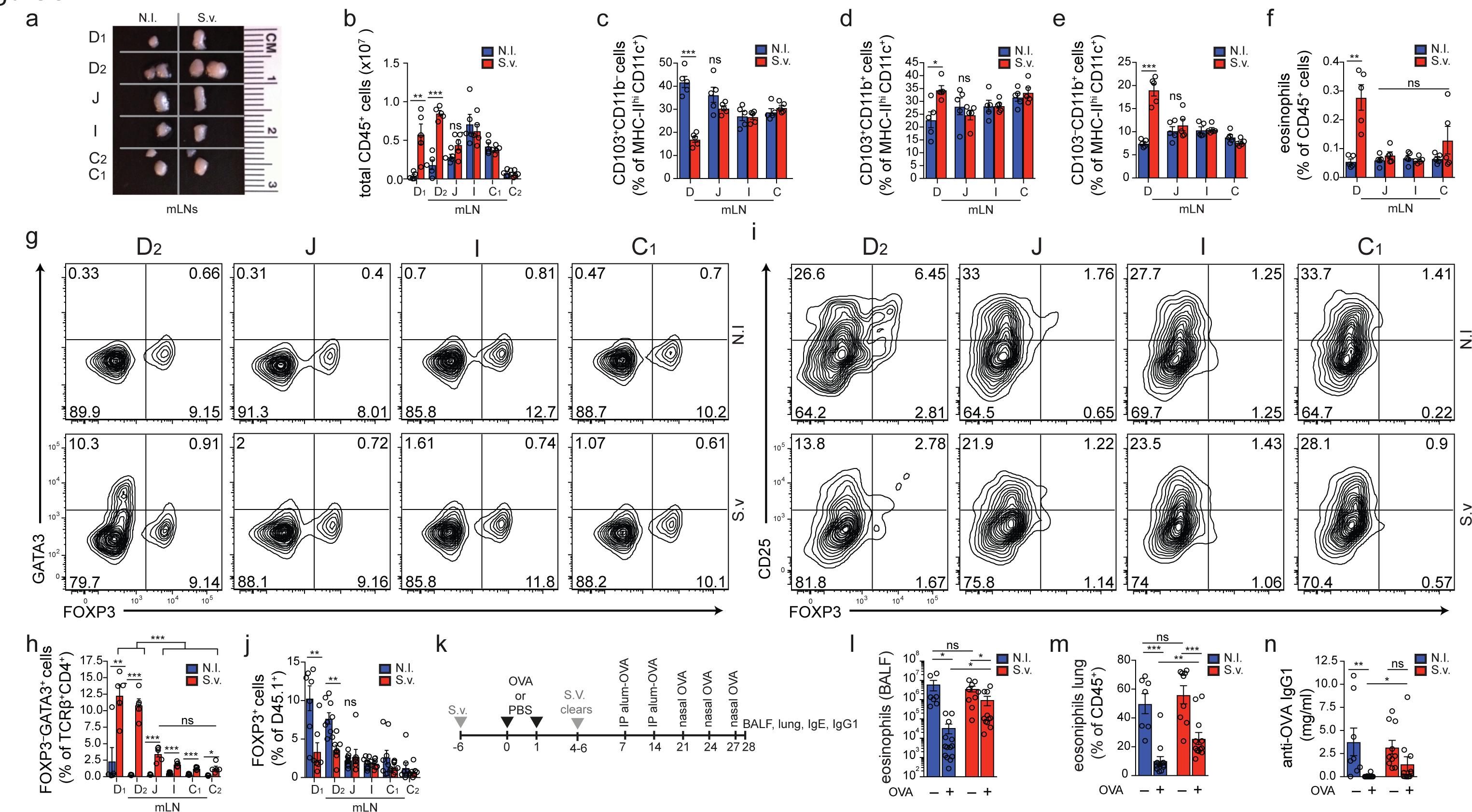
Duodenal infection leads to a compartmentalized immune conflict in the duodenal mLN and to compromised oral tolerance. **a**, Dissected indicated mLNs from non-infected (N.I.) mice or mice infected with 700 *S. venezuelensis* (S.v.) larvae 8 days prior to harvest. **b**, Total CD45^+^ cell counts (*n*=5). Data representative of 3 independent experiments. **c-f**, Frequency of CD103^+^CD11b^−^ (**c**), CD103^+^CD11b^+^ (**d**) and CD103^−^CD11b^+^ (**e**) among MHCII^hi^CD11c^+^ cells and eosinophils among CD45^+^ cells (**f**) (*n* = 5). Data representative of 2 independent experiments. **g**, Representative flow cytometry plot of GATA3^+^ and FOXP3^+^ CD4^+^ T cells. **h**, Frequency of GATA3^+^ CD4^+^ T cells (*n* = 5). Data representative of 3 independent experiments. **i**, Representative flow cytometry plot of FOXP3^+^ and CD25^+^ CD45. 1 OT-II cells in indicated mLNs 8 days after infection of mice with S.v. larvae or N.I. **j**, Frequency of total FOXP3^+^ among CD45.1^+^ cells in indicated mLNs 60 h post adoptive transfer of 1 x 10^6^ naïve CD45.1^+^ OT-II cells into CD45.2 hosts (*n*=8) infected with S.v. larvae or N.I. 8 days and gavaged with OVA 48 h and 24 h prior to analysis. Data pooled from two independent experiments. **k**, Scheme of oral tolerance experimental set-up in S.v. infected mice., **l**-**n**, Total eosinophils in bronchoalveolar lavage fluid (BALF) (**l**), frequency of eosinophils among CD45^+^ cells in lung tissue (**m**) and OVA-specific IgG1 levels in serum (n) from mice infected with S.v. or N.I during antigen feeding (+OVA groups) or no feeding (-OVA groups), at 21 d after first immunization with OVA-alum (**k**) (*n*=13 for +OVA groups, *n*=10 for -OVA groups). Data are pooled from two independent experiments. D=duodenum, whereby D1 portal LNs and D2 distal duodenum mLNs, J=jejunum, I=ileum, C1=cecal-colonic mLN, C2=ascending colonic mLN, C=colon. \**P* < 0.05, \*\**P* < 0.01 and \*\*\**P* < 0.005 (one-tailed *t*-test or ANOVA).

**Extended Figure 3.**
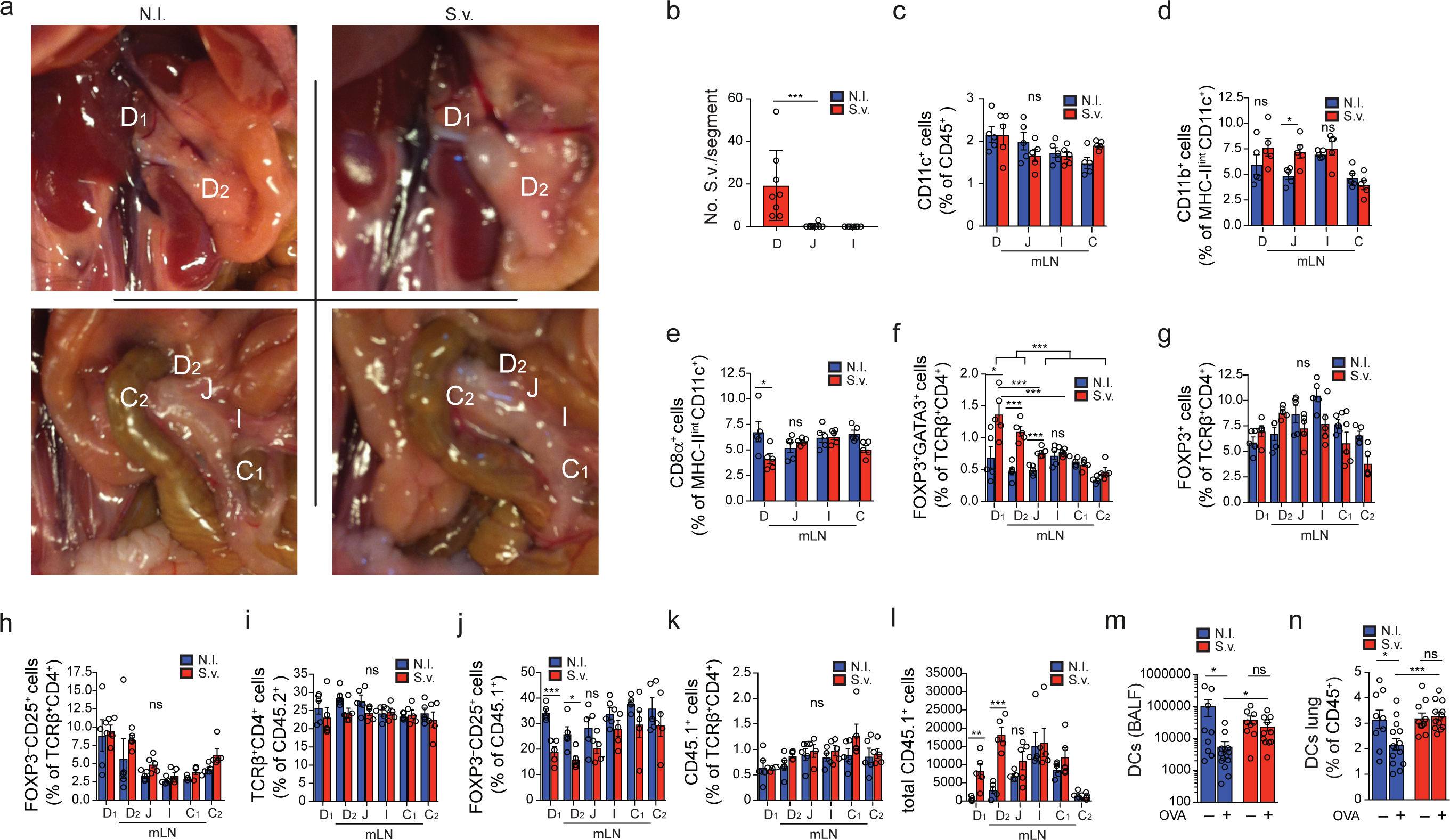
Extended analysis of the regional immune perturbation upon *S. venezuelensis* infection. **a**, Indicated mLNs positions in non-infected (N.I.) mice or mice infected with 700 *S. venezuelensis* (S.v.) larvae 8 days prior to harvest. **b**, Quantification of mature S.v. worms in gut epithelium 8 days after infection with 700 S.v. larvae. n=8, pooled from two independent experiments. **c**-**e**, Frequency of CD11c^+^ among CD45^+^ (**c**), and CD11b^+^(**d**) and CD8α^+^(**e**) among MHCII^int^CD11c^+^ cells (*n* = 5). Data representative of 2 similar experiments. **f**-**i**, Frequency of GATA3^+^ FOXP3^+^ (**f**), GATA3^−^ FOXP3^+^ (**g**), CD25^+^ FOXP3^−^ (**h**) among TCRβ^+^CD4^+^ cells and TCRβ^+^CD4^+^ among CD45^+^ (**i**) (*n* = 5). Data representative of three independent experiments. **j**-**l**, Frequency of total FOXP3^−^CD25^+^ among CD45.1^+^ (**j**) CD45.1^+^ among TCRβ^+^CD4^+^ (**k**) and total CD45.1+cells (**l**) in indicated mLNs 60 h post adoptive transfer of 1 x 10^6^ naïve CD45.1^+^ OT-II cells into CD45.2 hosts (*n*=4) infected with S.v. or N.I. 8 days and gavaged with OVA 48 h and 24 h prior to analysis. Data representative of two independent experiments. **m**, **n**, Total DCs in bronchoalveolar lavage fluid (BALF) (**m**), frequency of eosinophils among CD45^+^ cells in lung tissue (**n**) from mice infected with S.v. or N.I during antigen feeding (+OVA groups) or no feeding (-OVA groups), at 21 d after first immunization with OVA-alum (**k**) (*n*=13 for +OVA groups, *n*=10 for -OVA groups). Data are pooled from two independent experiments. D=duodenum, whereby D1 portal LNs and D2 distal duodenum mLNs, J=jejunum, I=ileum, C1=cecal-colonic mLN, C2=ascending colonic mLN, C=colon. \**P* < 0.05, \*\**P* < 0.01 and \*\*\**P* < 0.005 (one-tailed *t*-test or ANOVA).

We next investigated if the environmental context dictates T cell fate in the resistance-prone distal intestine-draining mLNs. We chose to track responses to Segmented Filamentous Bacteria (SFB), pathobionts that elicit a strong T_H_17 response and preferentially colonize the ileum and colon^27–29^. Monocolonization of GF mice with SFB led to the enrichment of RORγt^+^CD4^+^ T cells in the I-and C-mLNs 8 days after gavage (Fig. 4a). In the Peyer’s Patches (PP), lymphoid structures in the small intestine itself, the SFB colonization effect was less restricted, resulting in expansion of both RORγt^+^CD4^+^ T and IgA^+^CD95^+^ B cells in the LNs draining proximal small intestine regions (Fig. 4b, c). To monitor SFB responses in the context of a full microbiota we adoptively transferred naïve SFB-specific CD4^+^ T cells (7B8tg cells)^3^ into mice colonized with SFB 7 days before, and analysed their fate 60 h later. We only recovered significant numbers of transferred cells in the I-and C-mLNs (Fig. 4d-f, Extended Data Fig. 4a, b), and these effects were even more regionally-restricted in Taconic mice harbouring SFB through parental transmission (Extended Data Fig. 4c, d), suggesting that recent SFB colonization leads to transient, yet mild SFB dysbiosis in the duodenum and jejunum (Extended Data Fig. 4e, f). Because T_H_17 polarization was concentrated in LNs draining the distal segments of the intestine, we anticipated that surgical removal of the ileal and cecal mLNs (I and C1 in Fig. 1h) would abrogate SFB-specific T_H_17 responses. Removal of ileal-and cecal-mLNs did not result in evident lymphatic rerouting of antigen and migratory DCs to alternate mLNs, as indicated by fast green injection into the intestinal lymphatics (Extended Data Fig. 4g). However, distal mLN removal led to ectopic seeding by 7B8tg cells in D-, J-and ascending C-mLNs (C2 in Fig. 1h) and pronounced RORγt^+^ cell generation (Fig. 4d-f; in Taconic mice, Extended Data Fig. 4c, d). Distal mLN removal also resulted in a selective swelling of the ascending C-mLN, indicating that it partially took over the role of the cecal-colonic mLN (Extended Data Fig. 4h, i). This loss of compartmentalized SFB response was also reflected back in the gut tissue, since in ^ΔicLN mice SFB-specific T_H_17 cells were found in the duodenum and jejunum, while they were confined to the^ ileum and colon in sham-operated mice (Fig. 4g; Extended Data Fig. 4j-o, Taconic mice, Extended Data Fig. 4p-t), suggesting that in absence of the primary draining mLNs, T cells may instead traffic to mLNs draining the adjacent gut regions, where smaller amounts of cognate antigen are found (Extended Data Fig. 4e, f). These results indicate that the type and magnitude of immune responses can be preserved in the absence of segment-specific drainage. Nevertheless, disruption in lymphoid structures draining specific niches results in displacement of segment-specific adaptive immunity. Therefore, in addition to its role in properly distributing proximal and distal gut immune responses, compartmentalised drainage may be required for appropriate homing of T cells to specific gut segments.

**Figure 4.**
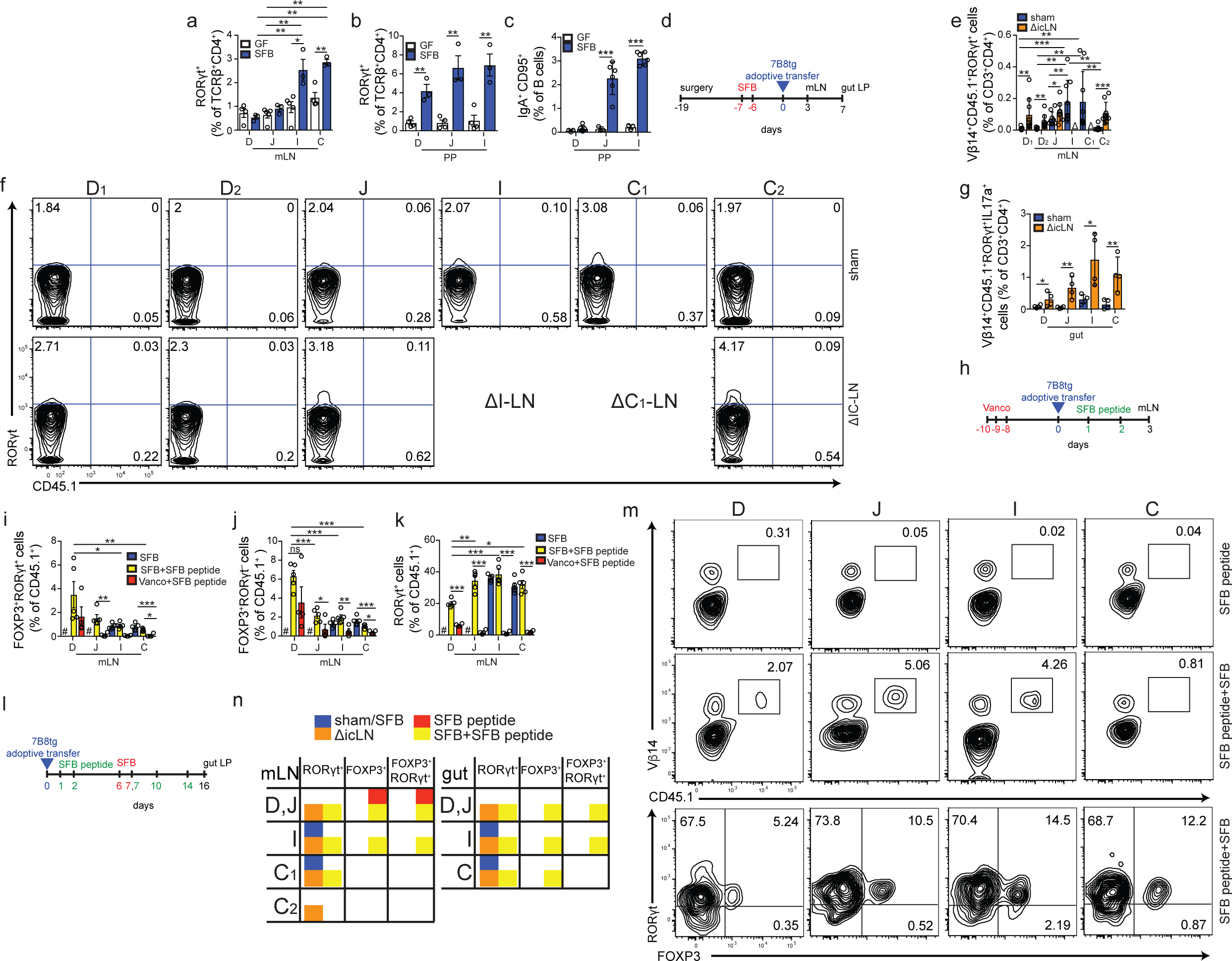
Response to an ileal pathobiont is delocalized and reprogrammed by perturbed antigen delivery to mLNs. **a**, **b**, Frequency of RORγt^+^ among TCRβ^+^CD4^+^ cells in indicated mLNs (**a**) or Peyer’s Patches (PP) from indicated intestinal segment (**b**) of GF mice or mice monocolonized with SFB 8 days prior to analysis (*n*=3). Data representative of two independent experiments. **c**, Frequency of IgA^+^CD95^+^ among B cells in PP from indicated intestinal segment of GF mice or mice monocolonized with SFB 4 weeks prior to analysis (*n*=3). Data representative of two independent experiments. **d**, Scheme of experimental design for **e**-**g**. **e**, Frequency of Vβ14^+^CD45.1^+^ RORγt^+^ cells among CD3^+^CD4^+^ cells in indicated mLN of mice with sham operation (*n*=12) or surgical removal of the I-and C1-mLN (ΔicLN, *n*=14) 60 h (day 3, **d**) after adoptive transfer of 4 x 10^5^ naïve SFB specific CD45.1^+^ 7B8tg cells. Graph represents pooled data from 3 independent experiments with *n*=4-5 per group each. **f**, Representative flow cytometry plots of RORγt^+^ and CD45.1^+^ frequency among CD3^+^CD4^+^ cells in indicated mLN of sham or ΔicLN mice as quantified in **e**. **g**, Frequency of Vβ14^+^CD45.1^+^ IL17a^+^ cells among CD3^+^CD4^+^ cells in lamina propria of indicated gut segment of sham or ΔicLN mice (*n*=4) 7 days after adoptive transfer of 5,000 naïve CD45.1^+^ 7B8tg cells. **h**, Scheme of experimental design for data **i**-**k**. **i**-**k**, Frequency of FOXP3^+^RORγt^+^ (**i**), FOXP3^+^RORγt^−^ (**j**) and FOXP3^+^RORγt^+^ (**k**) among Vβ14^+^CD45.1^+^ cells in indicated mLNs 60 h (day 3, **h**) after adoptive transfer of 4 x 10^5^ naïve CD45.1^+^ 7B8tg cells into untreated Taconic B6 hosts (SFB, *n*=4) or treated with SFB peptide (*n*=5) or Vanomycin and SFB peptide (*n*=4). Data representative of two similar experiments. **l**, Scheme of experimental design for data in **m**. **m**, Representative flow cytometry plots of Vβ14^+^ and CD45.1^+^ frequency among CD3^+^CD4^+^ cells (top rows) or FOXP3^+^and RORγt^+^ Vβ14^+^CD45.1^+^ cells (bottom row) in lamina propria of indicated gut segment of mice fed SFB peptide and subsequently colonized with SFB or not, 16 days after adoptive transfer of 5,000 naïve CD45.1^+^ 7B8tg cells as depicted in **l**. **n**, Summary table of 7B8tg CD4^+^ T cell fate in mLN and lamina propria upon mLN surgery and SFB peptide administration. D=duodenum, whereby D1 portal LNs and D2 distal duodenum mLNs, J=jejunum, I=ileum, C1=cecal-colonic mLN, C2=ascending colonic mLN, C=colon. \**P* < 0.05, \*\**P* < 0.01 and \*\*\**P* < 0.005 (one-tailed *t*-test or ANOVA).

**Extended Figure 4.**
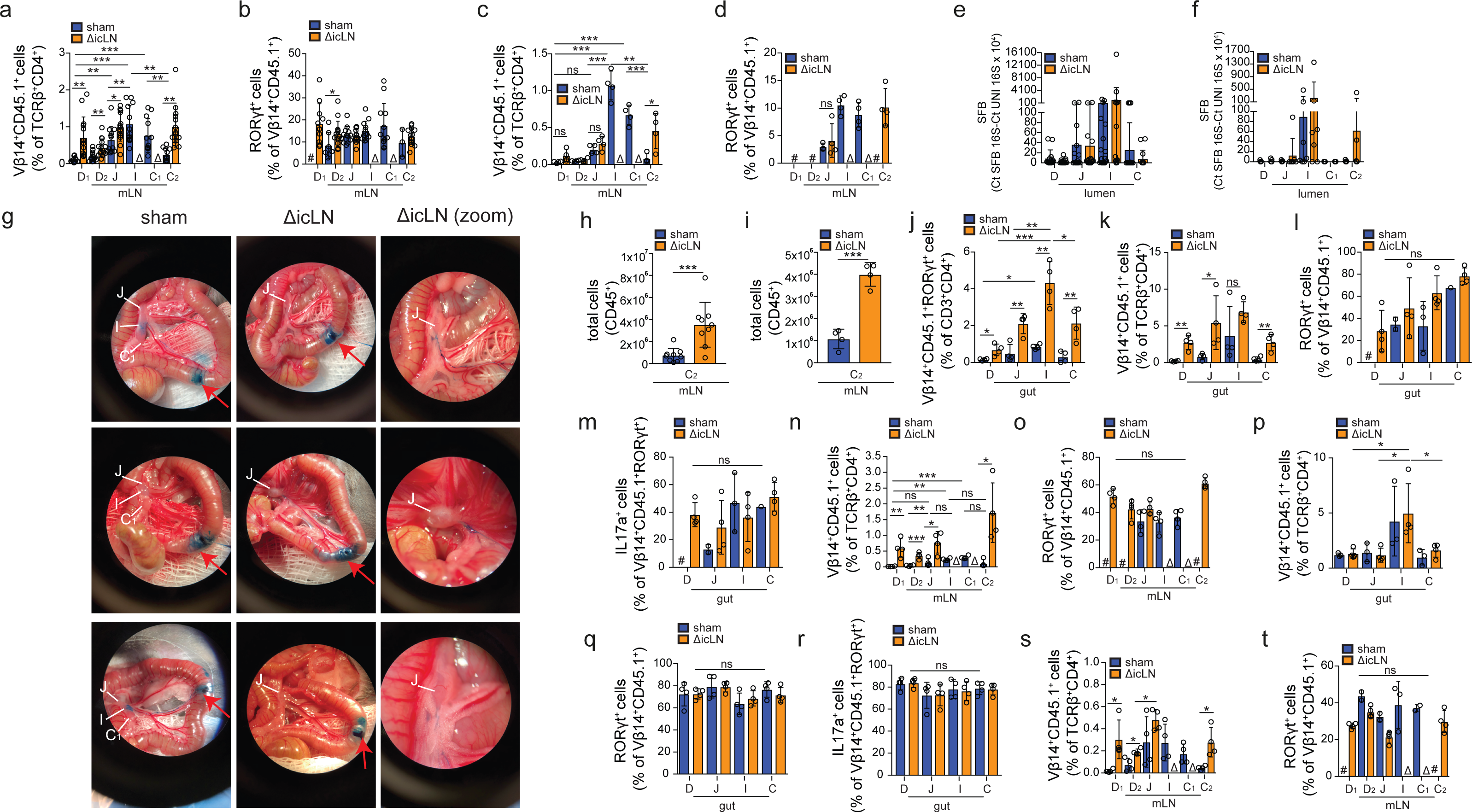
Extended characterization of the impact of distal lymph node surgery on the compartmentalized response to SFB. **a,b**, Frequency of Vβ14^+^CD45.1^+^ among CD3^+^CD4^+^ (**a**) and RORγt^+^ among Vβ14^+^CD45.1^+^ (**b**) cells in indicated mLN of mice with sham operation (*n*=12) or surgical removal of the I-and C1-mLN (ΔicLN, *n*=14) 60 h (day 3, **Fig. 4d**) after adoptive transfer of 4 x 10^5^ naïve SFB specific CD45.1^+^ 7B8tg cells. Graph represents pooled data from 3 independent experiments with *n*=4-5 per group each. **c**, **d**, Frequency of Vβ14^+^CD45.1^+^ among CD3^+^CD4^+^ (**c**) and RORγt^+^ among Vβ14^+^CD45.1^+^ (**d**) cells in indicated mLN of Taconic sham or ΔicLN mice (*n*=4) treated as **b**. **e**, **f**, Relative quantification of SFB specific 16S in luminal contents of indicated gut segment from recently SFB-colonized Jax B6 (**e**, *n*=13 per group from 4 independent experiments) or parentally-colonized Taconic (**f**, *n*=7 per group from 4 independent experiments) mice after sham or ΔicLN surgery. **g**, Picture of fast green tracing of ileal lymphatic drainage to the mLNs, 5 min after injection of 3 µl fast green in sham operated (left column, biological triplicates) or ΔicLN (middle and zoomed in right column, biological triplicates) mice 3 weeks after surgery. Red arrows indicate site of injection, jejunal (J), ileal (I) and cecal-colonic (C1) mLNs are indicated. **h**, **i**, Total CD45^+^ counts in ascending colonic mLN of sham or ΔicLN Jax (**g**, *n*=9 per group, pooled from 2 independent experiments) or Taconic (**i**, *n*=4, representative of two independent experiments) mice. **j**-**o**, Frequency of Vβ14^+^CD45.1^+^RORγt^+^ among CD3^+^CD4^+^ (**j**), Vβ14^+^CD45.1^+^ among CD3^+^CD4^+^ (**k**), RORγt^+^ among Vβ14^+^CD45.1^+^ (**l**) and IL17a^+^ among RORγt^+^Vβ14^+^CD45.1^+^(**m**) cells, or Vβ14^+^CD45.1^+^ among CD3^+^CD4^+^ (**n**), RORγt^+^ among Vβ14^+^CD45.1^+^ (**o**) in indicated lamina propria (**j**-**m**) or mLNs (**n**, **o**) of sham or ΔicLN Jax mice 7 days after adoptive transfer of 5000 naïve CD45.1^+^ 7B8tg cells (n=4). **p**-**t**, Frequency of Vβ14^+^CD45.1^+^ among CD3^+^CD4^+^ (**p**), RORγt^+^ among Vβ14^+^CD45.1^+^ (**q**) and IL17a^+^ among RORγt^+^Vβ14^+^CD45.1^+^(**r**) cells, or Vβ14^+^CD45.1^+^ among CD3^+^CD4^+^ (**s**), RORγt^+^ among Vβ14^+^CD45.1^+^ (**t**) in indicated lamina propria (**p**-**r**) or mLNs (**s**, **t**) of sham or ΔicLN Taconic mice 7 days after adoptive transfer of 5000 naïve CD45.1^+^ 7B8tg cells (n=4). # indicates fewer than 200 cells were recovered. D=duodenum, whereby D1 portal LNs and D2 distal duodenum mLNs, J=jejunum, I=ileum, C1=cecal-colonic mLN, C2=ascending colonic mLN, C=colon. \**P* < 0.05, \*\**P* < 0.01 and \*\*\**P* < 0.005 (one-tailed *t*-test or ANOVA).

We finally asked how the intestinal immune system handles an antigen delivered in multiple contexts, and what the consequences on tissue immunity will be. We delivered SFB as an oral antigen by gavaging its immune-dominant peptide, which is recognized by 7B8tg cells, to SFB-colonized Taconic mice and monitored 7B8tg seeding and fate in the mLNs 60 h later. SFB peptide gavage led to induction of RORγt^+^ and RORγt^−^ pTreg 7B8tg cells that followed the characteristic proximal to distal pTreg gradient observed upon dietary antigen administration, with or without SFB clearance by Vancomycin treatment (Fig. 4h-j, Extended Data Fig. 5a). In fact, these effects were preserved in ΔicLN mice, confirming a preeminent Treg-inducing capacity for the D-mLNs even during perturbation in the distal gut (Extended Data Fig. 5b-e). However, in SFB+ mice we also observed a substantial accumulation of RORγt^+^ 7B8tg cells in the D-and J-mLNs, while if SFB was cleared with Vancomycin we recovered very few RORγt^+^ cells in these mLNs (Fig. 4k). Conversely, priming by SFB peptide gavage prior to SFB colonization resulted in expansion of 7B8tg cells in the duodenum and jejunum in addition to the ileum, a significant fraction of which were RORγt^+^ pTreg cells (Fig. 4l, m, Extended Data Fig. 5f-j). These data show that adaptive immune responses towards distal challenges can influence proximal immune priming, suggesting a degree of plasticity in the compartmentalization of gut immunity, which may ensure a proper balance between resistance and tolerance (Fig. 4n). It remains to be determined whether specific DC subpopulations with varying tolerogenic or inflammatory roles^13^ along mLNs (Fig. 2) mediate these distinct effects towards soluble or pathogen-derived antigens.

**Extended Figure 5.**
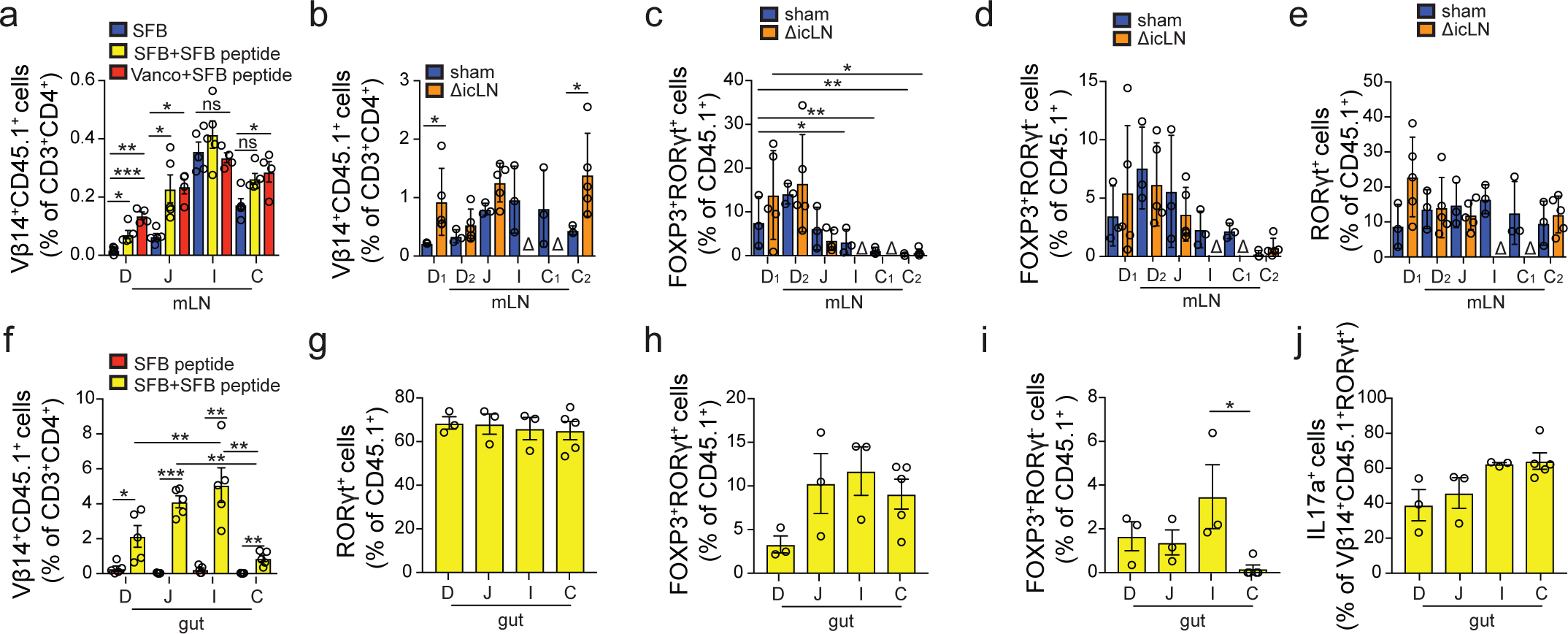
Extended analysis of the immune conflict imposed by SFB as an oral antigen. **a** Frequency of Vβ14^+^CD45.1^+^ among CD3^+^CD4^+^ cells in indicated mLNs 60 h (day 3, **Fig. 4h**) after adoptive transfer of 4 x 10^5^ naïve CD45.1^+^ 7B8tg cells into untreated Taconic B6 hosts (SFB, *n*=4) or treated with SFB peptide (*n*=5) or Vanomycin and SFB peptide (*n*=4). Data representative of two similar experiments. **b**-**e**, Frequency of Vβ14^+^CD45.1^+^ among CD3^+^CD4^+^ (**b**), FOXP3^+^RORγt^+^ (**c**), FOXP3^+^RORγt^−^ (**d**) and FOXP3^−^RORγt^+^(**e**) among Vβ14^+^CD45.1^+^ cells in indicated mLNs 60 h after adoptive transfer of 4 x 10^5^ naïve CD45.1^+^ 7B8tg cells into sham (*n*=3) or ΔicLN (*n*=5) Jax mice that were colonized with SFB 7 days prior to adoptive transfer and gavaged with 5 mg SFB peptide 48 h and 24 h prior to harvest. **f**, Frequency of Vβ14^+^CD45.1^+^ among CD3^+^CD4^+^ cells in lamina propria of indicated gut segment 16 days after adoptive transfer of 5000 naïve 7B8tg cells into mice fed SFB peptide alone (*n*=4) or that were subsequently colonized with SFB (*n*=4) (see **Fig. 4l**). **g**-**j**, Frequency of FOXP3^−^RORγt^+^ (**g**), FOXP3^+^RORγt^+^ (**h**), FOXP3^+^RORγt-(**i**), and IL17a^+^ among RORγt^+^Vβ14^+^CD45.1^+^(**j**) cells in lamina propria of indicated gut segment 16 days after adoptive transfer of 5000 naïve 7B8tg cells into mice fed SFB peptide and that were subsequently colonized with SFB (*n*=4, see **Fig. 4l**). D=duodenum, whereby D1 portal LNs and D2 distal duodenum mLNs, J=jejunum, I=ileum, C1=cecal-colonic mLN, C2=ascending colonic mLN, C=colon. \**P* < 0.05, \*\**P* < 0.01 and \*\*\**P* < 0.005 (one-tailed *t*-test or ANOVA).

Our data uncover a mechanism by which the intestinal immune system simultaneously handles regulatory versus pro-inflammatory responses: by anatomical segregation of these reactions into different, functionally specialized mLNs. The efficient drainage of dietary antigens into tolerance-promoting lymph nodes associated to the proximal intestine contributes to the understanding of food allergy prevention, and implicate duodenal infection and dysbiosis as environmental perturbations that can alter the allergic outcome. It is plausible that oral vaccination generally elicits diminished immune responses^30^ by this same mechanism. However, the tolerogenic duodenal route could be exploited therapeutically for inducing tolerance to otherwise inaccessible pro-inflammatory antigen sources such as dysbiotic ileal or colonic bacteria in inflammatory bowel diseases.

## Methods

### Mice

C57BL/6J CD45.1 (B6.SJL-Ptprca Pepcb/BoyJ) from Jackson laboratories, CD45.2 (C57BL/6) mice were purchased from the Jackson Laboratories or Taconic Farms, and CD45.1 OT-II TCR-transgenic were originally purchased from Taconic Farms and maintained in our facilities. B6.Cg-Tg(Itgax-Venus)1Mnz/J mice were provided by M. Nussenzweig (The Rockefeller University). C57BL/6-Tg (Tcra,Tcrb)2Litt/J (7B8tg) mice were either purchased from Jackson Laboratories or provided by D. Littman (NYU). Mice were used at 7-12 week old, and male mice were used throughout the study. Mice were maintained at the Rockefeller University animal facilities under specific pathogen-free conditions or germ free conditions. Germ-free C57BL/6 mice were generously provided by S. Mazmanian (Caltech) and imported into germ-free flexible film isolators (Class Biologically Clean Ltd.). Mice monocolonized with SFB were kept in flexible film isolators and originally colonized by gavage with fecal extract from SFB monocolonized mice kept at NYU (Littman lab). SFB colonization was verified by real time PCR using SFB-specific 16S primers; germ-free feces served as a negative, Taconic Farms B6 feces as a positive control. The mice were bred in our germ-free facility for maintenance of the strain and kept on sterilized Autoclavable Mouse Breeder Diet (5021, LabDiet, USA), which was also used for control (ex-germ free) SPF mouse breeding and maintenance.

### Rats

6-10 week old female Wistar rats were purchased from Charles River. Animal care and experimentation were consistent with the NIH guidelines and were approved by the Institutional Animal Care and Use Committee at the Rockefeller University.

### Reagents

The following reagents were purchased from Sigma: Benzyl ether (108014), Dichloromethane (270997), Fast Green FCF (F7252), Heparin (H3393), Omeprazole (O104), Pluronic L-81 (435430), Pyrantel Pamoate (P6210), Ovalbumin (grade VI, A2512; grade III A5378) and Vancomycin (V2002). LPS-free ovalbumin was from Hyglos, Germany (Cat. no 77161). ^3^H-retinol and Na^125^I were from Perkin-Elmer.

### Antibodies, staining and flow cytometry

Fluorescent-dye-conjugated antibodies were purchased from BD (USA) (anti-CD45.2, 560693; anti-CD103, 557495; anti-Ly6C, 560595; anti-NK1.1, 562921; anti-SiglecF, 552126; anti-RORγt, 562894; anti-IL17A, 56022 and 559502; anti-Vβ14, 553258), eBioscience (USA) (anti-B220, 48-0452-82; anti-CD3e, 48-0031-82; anti-CD4, 83-0042-42; anti-CD8α, 56-0081-82; anti-CD25, 17-0251-82; anti-CD11b, 47-0112-82; anti-CD11c, 25-0114-82, 17-0114-82 and 56-0114-82; anti-CD45, 25-0451-82; anti-CD45.1, 25-0453-82; anti-CD69; anti-FOXP3, 12-5773-82; anti-GATA3, 12-9966-42; anti-I-A/I-E (MHCII), 46-5321-82 and 56-5321-82; anti-Ly6G, 48-5931-82; anti-Vα2, 48-5812-82, and Streptavidin, 46-4317-82), or Biolegend (USA) (anti-CD8α, 100744; anti-CD11b, 101236; anti-CD64, 139306; anti-TCRβ, 109220). Additional antibodies were purchased from BioXCell and labelled in-house (anti CD4, BE0003-1; anti-CD8α, BE0004-1; anti-CD11b, BE007; anti-CD19, BE0150; anti-TCRβ, BE00102). Biotinylated antibodies were purchased from BD Pharmigen (anti-B220, 553086; anti-CD8α, 553029; anti-Cd11b, 553309; anti-Cd11c, 553800; anti-CD25, 553070; and anti-NK1.1, 553163; anti-TER-119, 553672;) or as follows: anti-IgG1, Bethyl, A90-105B; anti-IgG2c, Bethyl, A90-136B; anti-IgE, eBioscience, 13-5992-82; and anti-Neuropilin, R&D Systems, BAF566. Unconjugated antibodies used were anti-LYVE 1, R and D Systems AF2125; anti-GFP (Aves Labs, GFP-1020); anti-OVA IgG1, Biolegend, 520501; anti-IgE, Invitrogen, RMGE00; IgE isotype control, eBioscience 554118. Horseradish peroxidase conjugated Streptavidin was purchased from Jackson ImmunoResearch Laboratories, Inc. Aqua LIVE/DEAD^®^ Fixable Aqua Dead Cell Stain Kit, L-34965, Cell Trace CFSE and Violet Cell Proliferation kits (C34554 and C34557) were purchased from Life Technologies. For cytokine analysis in T cells, cells were incubated for 3.5 h in RPMI with 10% FBS, Brefeldin A (0.5 µg ml^−1^, Sigma B7651), ionomycin (0.5 µg ml^−1^, Sigma I0634), and phorbol 12-myristate 13acetate (100 ng ml^−1^, Sigma P8139), for cytokines in dendritic cells, cells were incubated for 6h in RPMI with 10% FBS and GolgiPlug (1 to 1000, BD Biosciences 555029). Cell populations were stained with Aqua in PBS, followed by incubation with Fc block and antibodies against the indicated cell surface markers in FACS buffer (PBS, 1% BSA, 10 mM EDTA, 0.02% sodium azide). The cells were analyzed live, or fixed in 1% PFA/PBS. For intracellular staining, cells were first stained for surface epitopes and then fixed, permeabilized and stained according to the manufacturer’s protocol (eBioscience 00-5123-43). Flow cytometry was performed on an LSRII (BD Biosciences) and analysed using FlowJo Software (Tree Star). Cell division index was calculated using the FlowJo formula (http://www.flowjo.com/v765/en/proliferation.html), whereby the index represents the fraction of total cell divisions over the calculated total starting cells.

### Tissue clearing, light sheet microscopy and image reconstruction

For tissue clearing and staining, the iDISCO protocol was followed as detailed on the continuously-updated website: http://idisco.info with the following specifics: Tissues were stained in primary and secondary antibodies for 4 days each, and were embedded in 1% agarose-TAE prior to the final dehydration. Blocks were imaged using a LaVision Ultramicroscope II and Software. Images were reconstructed using Imaris 8 Software.

### ^3^H-retinol biodistribution

Mice were fasted for 3h prior to gavage with 150 µl PBS with or without 5 µl chylomicron formation inhibiting Pluronic L-81, followed by 1 µCi ^3^H-retinol in 100 µl olive oil 30 minutes later, and sampled at indicated time points. Serum was sampled from submandibular vein (systemic) or from the portal vein. Lymph was collected from the thoracic duct under isoflurane anaesthesia using a custom-made glass needle (Micropipette puller, Sutter Instrument). Dissected organs were weighed prior to lysis by mechanical disruption in 0.5 ml hypertonic lysis buffer with 1% Triton-X 100, the lysate mixed with 7 ml Ultima Gold scintillation cocktail (Perkin Elmer) and accumulation of radioactivity measured on a scintillation counter. Input radioactivity was estimated by counting 10 % of the gavaged material.

### Segmentation of mesenteric lymph nodes

Mesenteric lymph nodes draining intestinal segments were determined anatomically by following the lymphatic vessels connecting the colon, ileum and jejunum to their lymph nodes. Duodenal lymph nodes were revealed by gavaging 100 µl of olive oil (Sigma) and determining the most stomach-proximal lymph nodes surrounded by chyle, indicative of duodenal drainage, 1h post gavage.

### Lymphocyte and APC isolation from lymph nodes

Tissues were dissected into cold HBSS, supplemented with Mg^2+^ and Ca^2+^, finely chopped and incubated in 400 U/ml Collagenase D (Roche) in HBSS for 25 min at 37°C, 5% CO_2_. Collagenase was quenched on ice by addition of final 10% FCS. Single cell suspensions were extracted from connective tissue by taking up and resuspending the digests five times. Erythrocytes were lysed by incubation in erythrocyte lysis buffer (Sigma) for 7 min at RT.

### Lymphocyte and APC isolation from small and large intestine

Intestines were separated from mesentery, and Peyer’s Patches (small intestine) and feces were removed. For segmentation of the small intestine, the upper 25% of the small intestine were taken as duodenum, the next 50% as jejunum and the last 25% as ileum. The cecum was included in the preparation of the large intestine. Intestines were cut longitudinally and washed twice in PBS. Tissue was cut into 1 cm pieces, mucus was removed by incubating the tissue for 10 min in PBS and 1 µM DTT, and the epithelium removed by two incubations in 25 ml of HBSS + 2% FCS + 30 mM EDTA for 10 min at 37°C at 230 rpm with vigorous shaking after each incubation. Tissues were washed in PBS over a sieve, then finely chopped and digested in 6 ml of RPMI per gut segment (Gibco), 2% FCS, 200 µg/ml DNaseI (Roche) and 2 mg/ml Collagenase 8 (Gibco) for 45 min at 37°C, 5% CO_2_. Digests were taken up and resuspended 10 times, passed through a sieve and the collagenase quenched by addition of 15 ml of cold RPMI, 2% FCS. Cell pellets were resuspended in 40% Percoll (BD Pharmigen) complemented with RPMI, 2% FCS, passed through a 100 µm mesh and separated by centrifugation in a discontinuous Percoll gradient (80%/40%) at 1000 *g* for 25 min at room temperature (RT). APCs and lymphocytes were isolated from the interphase, washed, and stained for FACS analysis or subjected to re-stimulation.

### Cell isolation for RNA-seq

Cells were sorted using a FACS Aria cell sorter flow cytometer (Becton Dickinson). MLN dendritic cells were pre-enriched using a Pan Dendritic Cell Isolation Kit (130-100-875, Miltenyi Biotec) and LS MACS Separation Columns (Miltenyi Biotec). 14 week old C57BL/6 males served as mLN donors, and biological triplicates or quadruplicates were collected. Dendritic cells were sorted as Aqua^−^CD45^+^Lin^−^(CD3^−^B220^−^ NK1.1^−^CD19^−^) CD11c^hi^, and the subpopulations further as MHCII^hi^CD103^+^CD11b^−^ and MHCII^hi^CD103^+^CD11b^+^. Three hundred cells were sorted directly into 25 µl TCL buffer (Qiagen, 1031576) supplemented with 1% β-mercaptoethanol at single cell precision. Samples were kept at room temperature for 5 min, spun down and kept at -80 **°**C until further processing.

### RNA-seq library preparation and sequencing

RNA was isolated using RNAClean XP beads (Agentcourt, A63987) on a magnetic stand (DynaMag, Invitrogen 12331D). Reverse transcription primers were: P1-RNA-TSO: Biot-rArArUrGrArUrArCrGrGrCrGrArCrCrArCrCrGrArUrNrNrNrNrNrNrGrGrG, P1-T31: Biot-AATGATACGGCGACCACCGATCG31T, P1-PCR: Biot-GAATGATACGGCGACCACCGAT. RNA was eluted for 1 min in RT-cDNA synthesis mix 1 (0. 5 µl P1-T31 (20uM), 0.3 µl RNasin plus (Promega, N2615), 1.5 µl 10 mM dNTP, 3.5 µl 10 mM Tris pH 7.5-0.5% IGEPAL CA-630 (Sigma) and 1.7 µl RNase free ddH_2_O) and the beads pipetted up and down ten times. The eluted sample was then incubated for 3 min at 72°C, followed by 1 min on ice, then 7.5 µl of mix 2 was added (3 µl 5X FS Buffer SS, 0.375 µl 100 mM DTT, 0.375 µl RNasin plus, 0.5 µl P1-RNA-TSO (40uM), 0.75 µl Maxima RT Minus H (Thermo Scientific, EP0751), 1.8 µl 5M Betaine (Sigma, B0300), 0.9 µl 50 mM MgCl2 and 0.175 µl RNase free ddH_2_O. Reverse transcription (RT) occurred during a thermal cycle of one cycle (90 min at 42 **°**C), 10 cycles (2 min at 50°C, 2 min at 42°C) and one cycle (15 min 70 °C), and the product was kept at 4 °C. The cDNA was then amplified using 15 µl of the RT product, 20 µl 2x KAPA HiFi HS Ready Mix (Kapabiosystems, KK2601), 1.5 µl P1-PCR (10uM), and 3.5 µl RNase free ddH_2_O. Amplification occurred during following cycle: One cycle of 3 min at 98 °C, 20 cycles (15 sec at 98 °C, 20 sec at 67 °C, 6 min at 72 °C), one cycle (5 min at 72 °C), and the product was kept at 4 °C. 20 µl of PCR product were cleaned up using 16 µl RNAClean XP beads. The cDNA was eluted in 20 µl RNase free ddH_2_O and kept at -20 °C. Isolated amplicons were confirmed to be 1500-2000 bp long by a High Sensitivity DNA Assay (Bioanalyzer). Concentration of all sample was measured on a Qubit fluorometer (Thermo Fisher), all samples were adjusted to 0.1 ng/µl with ddH_2_O, and 2.5 µl cDNA were subjected to Nextera XT DNA Library preparation (Illumina) using a Nextera XT Index Kit (Illumina, FC-131-1002) and according to the manufacturer’s protocol, except that all volumes were used at 0.5 x of the indicated volumes. Sample quality was again verified by Bioanalyzer, sample concentrations measured on the Qubit fluorometer and adjusted to a concentration of 4.54 ng/ µl. All samples were pooled at equal contribution and run in multiple lanes. Sequencing was performed using 75 base single-end reading on a Nextseq instrument (Illumina).

### RNA-seq data analysis

Equal contribution of samples to reads was verified. Raw reads were aligned using Kallisto and differential expression was performed with DESeq2. A p-value of 0.05 was used as a cutoff to determine differentially expressed genes highlighted in volcano plots. PCA plots and volcano plots were generated in R. A ranked file was generated based on log2 fold changes and subsequently run as a GSEA Preranked analysis (GSEA v3.0) with the c5.all.v6.1.symbols.gmt (Gene Ontology) gene set database. All pathways with a false detection rate (FDR) of 0.25 or less were considered significantly different between samples.

### RALDH activity assay

RALDH activity was determined using the Aldefluor kit (STEMCELL^TM^ Technologies) according to the manufacturer’s protocol.

### ^125^I-OVA labeling and biodistribution

Iodination of OVA was performed as described previously^14^. Mice were fasted for 3 h prior to gavage with 4 x 10^6^ CPM ^125^I-OVA and 50 mg cold OVA (grade III) in 200 µl PBS, and samples were taken at 1 h and 5 h post gavage. Wet weight of tissues was taken prior to measuring radioactivity on a gamma counter (Packard Cobra). Input radioactivity was estimated by counting 10 % of the gavaged material.

### Adoptive T cell transfer

Naïve CD4 T cells from spleen and lymph nodes were isolated by negative selection using biotinylated antibodies against CD8α, CD25, CD11c, CD11b, TER-119, NK1.1, and B220 and anti-biotin MACS beads (Miltenyi Biotec). Purity of transgenic CD4^+^ T cells was verified by flow cytometry (CD45.1^+^Vα2^+^Vβ5^+^CD25^−^ for OT-II cells, CD45.1 CD45.1^+^ Vβ14^+^CD25^−^ for 7B8tg cells, typically >90%). T cells were labeled using the Cell Trace^TM^ Violet or CFSE Cell Proliferation Kit (Life Technologies). For OT-II cells, 2×10^6^ were transferred by retro-orbital injection under isoflurane gas anesthesia. For 7B8tg cells, 4 x 10^5^ cells were transferred for analysis of mLNs 60 h post-transfer and 5000 cells for analysis 7 days or more after transfer in the gut and mLNs.

### Oral antigen administration

OVA (grade III, Sigma, A5378) was administered at 50 mg in 200 µl PBS by oral gavage using metal gavage needles. Two doses were given with a 24 h interval. SFB peptide FSGAVPNKTD (custom made with N-terminal acetylation, LifeTein, LLC.) was administered as either 1 mg or 5 mg in 200 µl PBS by oral gavage after i.p. treatment with 1 mg proton pump inhibitor Omeprazole (in 200 µl PBS, 1 % Tween 80) given 24 h and 15 min prior to gavage to reduce digestion of peptide in the stomach. Control mice were only given Omeprazole.

### *S. venezuelensis* passage and infection

*S.venezuelensis* was maintained in Wistar rats by subcutaneous infection with 30,000 larvae. Day 6-8 p.i. the cecum containing eggs was harvested and spread on Whatman paper which was placed into a beaker with water at 28 °C. The hatching larvae were collected over 4 days and the cycle re-initiated. Mice were infected subcutaneously with 700 larvae/ mouse. Adult worm load was assessed in total epithelial scrapes of the gut.

### Alum immunization and airway challenge

Seven days after oral OVA administration, 4 µg of endotoxin-free OVA antigen adsorbed to 40 µl Imject™ Alum Adjuvant (Fisher Scientific) was injected i.p. in a final volume of 400 µl made up with PBS. Immunization was repeated after 7 days. To induce airway inflammation, mice were anesthetized and intranasally administered 10 µg of sterile OVA grade VI in 50 µl PBS (25 µl per nostril) on days 14, 17 and 21 after the first i.p. immunization. Total IgE was measured to confirm previous infection with *S. venezuelensis* (250-350 ng/ml in plasma compared to 5-10 ng in plasma of uninfected mice) ^14^.

### Bronchoalveolar Lavage (BAL), lung histology and infiltrate analysis by flow cytometry

Mice were anesthetized by i.p. injection of 0.35 ml of 2.5% avertin (Sigma), the trachea was cannulated and lungs were lavaged once with 0.5 ml and then 1.0 ml PBS. Total BAL cells were counted after erythrocyte lysis and stained for FACS analysis. Lungs were perfused via the right ventricle with 10 ml saline to wash out residual blood. One lobe was digested in 400 U/ml collagenase D/HBSS and processed for FACS analysis. Eosinophils were determined as CD45^+^SSA^hi^MHCII^−^CD11b^+^Ly6G^int^SiglecF^+^, neutrophils as CD45^+^SSA^hi^MHCII^−^CD11b^+^Ly6G^hi^SiglecF^−^, and DCs as CD45^+^MHCII^+^CD11c^+^CD64^−^SiglecF^−^.

### Anti-OVA IgG1 and total IgE ELISAs

ELISAS were performed as described previously^14^.

### SFB colonization and depletion

SFB was obtained from frozen stocks of cecal contents (kept at -80 °C for less than 6 months) of mice monocolonized with SFB, which were diluted in PBS (2 ml/cecum) and passed through a 70 µm mesh. Mice were colonized by two gavages of 0.4 ml of cecal content preparation, 24 h apart. For depleting SFB, mice were treated with 200 µl of 10 mg/ml Vancomycin 10, 9, and 8 days prior to adoptive transfer of 7B8 cells, and the cages were exchanged every time to minimize re-colonization by remnant SFB in feces. Colonization or depletion was verified by real time PCR of fecal DNA (Quick-DNA Fecal/Soil microbe miniprep kit, Zymo research, Cat. No. D6010). PCR was performed in presence of Power SYBR Green PCR master mix (Applied Biosystems, 4367659) on a Quant Studio 3 PCR machine (Applied Biosystems), using SFB specific 16S primers (fwd: GACGCTGAGGCATGAGAGCAT, rev: GACGGCACGGATTGTTATTCA) The SFB Ct value was normalized by the Ct obtained in the PCR using universal bacterial 16S primers (fwd: ACTCCTACGGGAGGCAGCAGT, rev: ATTACCGCGGCTGCTGGC).

### Lymph node surgery

Mice were anesthetized subcutaneously with 100 mg/kg ketamine (controlled substance provided by the Rockefeller University animal facility), 10 mg/kg xylazine (Akorn, inc.), and in presence of 5 mg/kg analgesic Meloxicam (Putney, Inc.) in 0.5 ml saline. The abdominal area was then shaved, and sterilized by three cycles of wiping with iodine solution and 70 % ethanol after 20 µl of 0.25% bupivacaine (Hospira, Inc.) was injected intra-dermally at the prospective site of incision. The mouse was placed on a heat mat and covered by a sterile surgical plastic with an opening above the abdomen. All work from here was performed aseptically. Skin was incised in the middle of the abdomen along the anterior-posterior axis of the mouse, and the peritoneum was cut along the linea alba. To expose the ileal and cecal lymph nodes, the cecum was gently pulled out using cotton tipped applicators soaked in saline. Lymph nodes were removed by holding onto the lymph node with tweezers and gently pulling it while slowly and closely cutting around the node with microsurgical scissors, such that no bleeding occurred. For sham operation the cecum was pulled out and the lymph nodes exposed for 2 min. The cecum was then placed back into its original position and the peritoneal cavity filled with 0.5 ml pre-warmed saline. The peritoneal muscles were aligned and sutured using absorbable suture (PDS*II, Ethicon), the skin closed with autoclips (Ken Scientific Corp.), and the wound covered with Triple Antibiotic Ointment (Honeywell Safety Products, USA). Mice were allowed to fully awake in a cage placed on a heat mat. The following day mice were monitored for agility and passing stool. Mice were treated with 0.3 mg/kg of buprinex (controlled substance) and 5 mg/kg Meloxicam 24 h after first Meloxicam and for another 3 days every 24 h.

### Fast green tracing

Mice were anesthetized with isoflurane and their peritoneal cavity exposed. 3-5 µl of 10 % Fast Green in PBS were injected into the muscularis of the terminal ileum using a nano-injection device (Nanoject III, Drummond). Spreading of green colour through the lymphatics was followed until it accumulated in the ileum draining lymph node and until it faded again (sham operated mice) and reached the duodenal lumen via the bile duct, no longer than 15 minutes. The same timing was applied in mice in which ileal and cecal mLNs had been removed, even though the dye never accumulated in a lymph node.

### Statistical analysis

Statistical analysis was performed in GraphPad Prism 7.0 software. Multivariate data was analyzed by applying one-way ANOVA and Tukey’s multiple comparison *post hoc* test, comparison between two treatment conditions by one-tailed unpaired Student’s t-test.. A *P* value of less than 0.05 was considered significant.

## Acknowledgements

We thank all Mucida Lab members past and present for assistance in experiments, in particular intestinal lymphocyte preparations, fruitful discussions and critical reading of the manuscript, Aneta Rogoz for the maintenance of gnotobiotic mice, Sara Gonzalez for maintenance of SPF mice, Tomiko Rendon and Beatriz Lopez for genotyping, Kristie Gordon and Kalsang Chhosphel for maintenance of the flow cytometers and assistance with sorting, the Rockefeller University Bio-imaging Research Center for assistance with the light sheet microscopy and image analysis, the Rockefeller University Genomics Center for RNA sequencing and the Rockefeller University employees for continuous assistance. We also thank to Michel Nussenzweig, Gabriel Victora (Rockefeller University) and Juan Lafaille (NYU) and their respective lab members for fruitful discussions and suggestions. We are also indebted to Dan Littman and Mo Xu (NYU) for generously providing 7B8tg mice and feces from SFB monocolonized mice; and Stephen Galli and Kazufumi Matsushita (Stanford University) for providing *S. venezuelensis* and guidance on how to maintain it. This work was supported by a Swiss National Science Foundation postdoctoral fellowship (D.E.); a CAPES fellowship (M.C.C.C.); the Leona M. and Harry B. Helmsley Charitable Trust, the Crohn’s & Colitis Foundation of America Senior Research Award, the Burroughs Wellcome Fund PATH Award, National Institute of Health grants R21AI31188, R01DK113375 and R01DK093674 (D.M.).

## Author contributions

D.E. initiated, designed, performed and analysed the research, and wrote the paper. M.C.C.C. designed and performed infection and surgery studies. L.M. performed the DC sorting and guided the RNA-seq library preparation, P.A.M. performed RNA-seq data alignment and analyses and fast green injection. A.L. performed radioactive studies. A.M.C.F initiated the *S. venezuelensis* model and contributed to the experimental design using the model. All authors edited the paper. D.M. initiated, designed and supervised the research, and wrote the paper.

## Materials & Correspondence

Daria Esterházy or Daniel Mucida

## Supplementary Video Legends

**Supplementary Video 1.** Duodenal lymphatics (LYVE-1, white) of 9 weeks old SPF mouse. Antibody stained solvent cleared tissue followed by light sheet microscopy and 3D software reconstruction.

**Supplementary Video 2.** Duodenal lymphatics (LYVE-1, white) with APCs (CD11c-GFP, red) of 7 weeks old SPF mouse. Antibody stained solvent cleared tissue followed by light sheet microscopy and 3D software reconstruction.

**Supplementary Video 3.** Duodenal and mLN lymphatics (LYVE-1, white) 9 weeks old SPF mouse. Antibody stained solvent cleared tissue followed by light sheet microscopy and 3D software reconstruction.

**Supplementary Video 4.** mLN chain to duodenum lymphatics (LYVE-1, white) of 8 weeks old SPS mouse. Antibody stained solvent cleared tissue followed by light sheet microscopy and 3D software reconstruction.

**Supplementary Video 5.** ileal and mLN lymphatics (LYVE-1, white) of 8 weeks old SPS mouse. Antibody stained solvent cleared tissue followed by light sheet microscopy and 3D software reconstruction.

**Supplementary Video 6.** mLN lymphatics (LYVE-1, white) with APCs (CD11c-GFP, red) of 8 weeks old SPS mouse. Antibody stained solvent cleared tissue followed by light sheet microscopy and 3D software reconstruction.

**Supplementary Video 7.** mLN chain lymphatics (LYVE-1, white) with APCs (CD11c-GFP, red) and insulin (green) of 9 weeks old SPS mouse. Antibody stained solvent cleared tissue followed by light sheet microscopy and 3D software reconstruction.

## References

1 Belkaid, Y. & Hand, T. W. Role of the microbiota in immunity and inflammation. Cell 157, 121–141, doi:10.1016/j.cell.2014.03.011 (2014).

2 Honda, K. & Littman, D. R. The microbiota in adaptive immune homeostasis and disease. Nature 535, 75–84, doi:10.1038/nature18848 (2016).

3 Yang, Y. et al. Focused specificity of intestinal T_H_17 cells towards commensal bacterial antigens. Nature 510, 152–156, doi:10.1038/nature13279 (2014).

4 Xu, M. et al. c-MAF-dependent regulatory T cells mediate immunological tolerance to a gut pathobiont. Nature 554, 373–377, doi:10.1038/nature25500 (2018).

5 Mucida, D. et al. Oral tolerance in the absence of naturally occurring Tregs. The Journal of clinical investigation 115, 1923–1933 (2005).

6 Randolph, G. J., Ivanov, S., Zinselmeyer, B. H. & Scallan, J. P. The Lymphatic System: Integral Roles in Immunity. Annual review of immunology 35, 31–52, doi:10.1146/annurev-immunol-041015-055354 (2017).

7 Fonseca, D. M. et al. Microbiota-Dependent Sequelae of Acute Infection Compromise Tissue-Specific Immunity. Cell 163, 354–366, doi:10.1016/j.cell.2015.08.030 (2015).

8 Cording, S. et al. The intestinal micro-environment imprints stromal cells to promote efficient Treg induction in gut-draining lymph nodes. Mucosal immunology 7, 359–368, doi:10.1038/mi.2013.54 (2014).

9 Faria, A. M. C., Reis, B. S. & Mucida, D. Tissue adaptation: Implications for gut immunity and tolerance. J Exp Med 214, 1211–1226, doi:10.1084/jem.20162014 (2017).

10 Bernier-Latmani, J. & Petrova, T. V. Intestinal lymphatic vasculature: structure, mechanisms and functions. Nature reviews. Gastroenterology & hepatology 14, 510–526, doi:10.1038/nrgastro.2017.79 (2017).

11 Hooper, L. V. & Gordon, J. I. Commensal host-bacterial relationships in the gut. Science 292, 1115–1118 (2001).

12 Mucida, D. et al. Reciprocal T_H_17 and regulatory T cell differentiation mediated by retinoic acid. Science 317, 256–260 (2007).

13 Durai, V. & Murphy, K. M. Functions of Murine Dendritic Cells. Immunity 45, 719–736, doi:10.1016/j.immuni.2016.10.010 (2016).

14 Esterhazy, D. et al. Classical dendritic cells are required for dietary antigen-mediated induction of peripheral Treg cells and tolerance. Nature immunology 17, 545–555, doi:10.1038/ni.3408 (2016).

15 Gobert, M. et al. Regulatory T cells recruited through CCL22/CCR4 are selectively activated in lymphoid infiltrates surrounding primary breast tumors and lead to an adverse clinical outcome. Cancer Res 69, 2000–2009, doi:10.1158/0008-5472.CAN-08-2360 (2009).

16 Houston, S. A. et al. The lymph nodes draining the small intestine and colon are anatomically separate and immunologically distinct. Mucosal immunology 9, 468–478, doi:10.1038/mi.2015.77 (2016).

17 Atarashi, K. et al. Induction of colonic regulatory T cells by indigenous Clostridium species. Science 331, 337–341, doi:10.1126/science.1198469 (2011).

18 Kim, K. S. et al. Dietary antigens limit mucosal immunity by inducing regulatory T cells in the small intestine. Science, doi:10.1126/science.aac5560 (2016).

19 Hadis, U. et al. Intestinal tolerance requires gut homing and expansion of FoxP3+ regulatory T cells in the lamina propria. Immunity 34, 237–246, doi:10.1016/j.immuni.2011.01.016 (2011).

20 Balmer, M. L. et al. The liver may act as a firewall mediating mutualism between the host and its gut commensal microbiota. Sci Transl Med 6, 237ra266, doi:10.1126/scitranslmed.3008618 (2014).

21 Bouziat, R. et al. Reovirus infection triggers inflammatory responses to dietary antigens and development of celiac disease. Science 356, 44–50, doi:10.1126/science.aah5298 (2017).

22 Mukai, K., Karasuyama, H., Kabashima, K., Kubo, M. & Galli, S. J. Differences in the Importance of Mast Cells, Basophils, IgE, and IgG versus That of CD4(+) T Cells and ILC2 Cells in Primary and Secondary Immunity to Strongyloides venezuelensis. Infection and immunity 85, doi:10.1128/IAI.00053-17 (2017).

23 Tussiwand, R. et al. Klf4 expression in conventional dendritic cells is required for T helper 2 cell responses. Immunity 42, 916–928, doi:10.1016/j.immuni.2015.04.017 (2015).

24 Wilson, M. S. et al. Suppression of allergic airway inflammation by helminth-induced regulatory T cells. J Exp Med 202, 1199–1212, doi:10.1084/jem.20042572 (2005).

25 Maizels, R. M. & McSorley, H. J. Regulation of the host immune system by helminth parasites. The Journal of allergy and clinical immunology 138, 666–675, doi:10.1016/j.jaci.2016.07.007 (2016).

26 Curotto de Lafaille, M. A. et al. Adaptive Foxp3+ regulatory T cell-dependent and -independent control of allergic inflammation. Immunity 29, 114–126, doi:10.1016/j.immuni.2008.05.010 (2008).

27 Sano, T. et al. An IL-23R/IL-22 Circuit Regulates Epithelial Serum Amyloid A to Promote Local Effector Th17 Responses. Cell 163, 381–393, doi:10.1016/j.cell.2015.08.061 (2015).

28 Atarashi, K. et al. T_H_17 Cell Induction by Adhesion of Microbes to Intestinal Epithelial Cells. Cell 163, 367–380, doi:10.1016/j.cell.2015.08.058 (2015).

29 Ivanov, I. I. et al. Induction of intestinal Th17 cells by segmented filamentous bacteria. Cell 139, 485–498, doi:10.1016/j.cell.2009.09.033 (2009).

30 Levine, M. M. Immunogenicity and efficacy of oral vaccines in developing countries: lessons from a live cholera vaccine. BMC Biol 8, 129, doi:10.1186/1741-7007-8-129 (2010).

